# The role of superior colliculus in a logistic decision

**DOI:** 10.64898/2026.06.05.730072

**Authors:** Flóra Takács, Célian Bimbard, George M. Booth, Magdalena Robacha, Maxwell Shinn, Karolina Z. Socha, Kenneth D. Harris, Philip Coen, Matteo Carandini

**Affiliations:** UCL Institute of Ophthalmology, University College London, London, UK; Sainsbury Wellcome Centre, University College London, London, UK; UCL Queen Square Institute of Neurology, University College London, London, UK; Department of Cell and Developmental Biology, University College London, London, UK; Francis Crick Institute, London, UK; David Geffen School of Medicine, University of California, Los Angeles, USA

## Abstract

In many circumstances the brain makes decisions according to logistic classification: summing weighted factors in favor of each option and choosing an option according to a softmax function. Here we find that the superior colliculus (SC) contributes additively to this computation, promoting competing options independently of stimuli. We trained mice to locate an audiovisual stimulus on their left or right, and found that SC encodes the auditory, visual, and action signals in largely separate neurons in different layers. Unilateral SC inactivation did not affect sensory sensitivity but reduced contralateral actions, mostly in favor of inaction. The effects of bilateral inactivation were additive, with balanced inactivations restoring left-right balance. A simple 3-option logistic classification model captured the results, with sensory and prefrontal cortex processing sensory signals and each side of SC additively promoting the contralateral action while slightly opposing the ipsilateral action. The SC thus simultaneously promotes all competing eligible actions.

## Introduction

To choose an appropriate action, the brain must often integrate diverse sensory and nonsensory signals. Across a broad range of tasks, this integration follows a simple computation: logistic classification^1,2^. In logistic classification, the evidence in favor of each action is a weighted sum of factors and the probability of each action results from a softmax function (which is simply a logistic function if there are only two options). Logistic classification is standard in machine learning^3^ and in economics, where it describes collective consumer choices^4–6^. In individual observers, it characterizes choices based on multiple factors^2^ (alone or in combination) such as expected value^1,7–9^, recent history^10,11^, and sensory stimulation^9,12,13^.

Logistic classification can also describe the integration of stimuli across modalities and the role of cortical areas in this integration^2,14^. When mice decide between left vs. right based on combinations of visual and auditory cues, they weigh and sum the visual and auditory signals and decide according to logistic classification^14^. Inactivating one side of the visual cortex affects only the visual term in the sum, reducing the contralateral visual sensitivity^14–16^. Similarly, inactivating the secondary auditory cortex affects only the auditory term, reducing the contralateral auditory sensitivity^14^. Finally, inactivating the prefrontal cortex affects both the visual and auditory terms, implicating this region in audiovisual integration^14^.

This description of audiovisual decisions as entirely cortical, however, is puzzling given the prominent role that could in principle be played by a subcortical structure: the superior colliculus (SC). First, the SC could contribute to audiovisual decision by processing auditory and visual inputs and even by integrating them ^17^. The superficial and intermediate SC layers contain visual and auditory maps ^17–20^, and neurons in intermediate layers have been reported to respond to both visual and auditory inputs, often with nonlinear interactions between the two ^21–31^. Second, the SC might contribute by eliciting actions ^32,33^. The intermediate and deep layers of SC are denoted as motor (SCm) because their stimulation can elicit diverse orienting movements ^32–35^ through projections to brainstem and spinal cord. Indeed, unilateral SC inactivation or ablation reduces contralateral orienting movements ^36–41^ (often favoring ipsilateral ones ^36,40–45^). The SCm might thus execute movements selected in striatum or cortex ^43,45–53^. Third, the SC might contribute a more cognitive role ^54,55^. The activity of SCm typically encodes spatial targets ^56–65^ rather than movement coordinates ^66–71^, suggesting contributions to a priority map ^72,73^. During perceptual decisions, the SCm can modulate stimulus sensitivity ^74–76^ and decision bias ^40,41,70,73,76–79^ perhaps through ascending thalamic projections ^80–82^ that in turn affect the cortex.

However, the causal and computational roles of SC in audiovisual integration and decision making remain unclear. First, there is debate as to the extent of audiovisual interactions in SC neurons ^83^,^84^ and there is limited^85^ evidence that the SC causally affects audiovisual behavior. Second, the SC might not be necessary for motor execution: bilateral SC inactivations or ablations seemingly restore the effects of unilateral manipulations (the “Sprague effect”) ^36,39,86–90^. Third, there is debate as to the possible cognitive roles of SC in influencing sensorimotor target selection and perceptual decisions: many apparently complex effects of SC on perceptual decisions^79,91–94^ might be parsimoniously explained by an effect on decision bias ^95^.

To define this causal and computational role of SC, we used recordings and inactivations in mice performing an audiovisual localization task and leveraged a simple yet powerful logistic model of the decision. Our results reveal that the mouse SC plays a very different role than the cortex in audiovisual decisions. The SC encodes vision, audition, and action independently, and does not contribute to audiovisual integration. Rather, it contributes additive factors to the logistic process, which promotes all eligible actions independently of sensory signals. We call these factors action pressures, as they advantage eligible actions over ineligible actions, which include inaction. The action pressures oppose each other and largely cancel out in the overall logistic competition, but they vary within and across trials, contributing to the variability of choices.

## Results

We trained mice to turn a steering wheel to report the location of a stimulus ^11,15^ defined by visual and auditory cues^14^ (**Figure 1**a). The auditory cues (pink noise pulsing at 8 Hz) were presented from one of 3 loudspeakers located to the left (−60°), in front (0°), or to the right (+60°). The visual cues (checkerboards flashing at 8 Hz) appeared on the left or on the right (±60°), and varied by contrast. Unrewarded trials were followed by a 1.5 s timeout. Failure to respond within 1.5 s was classified as a NoGo (**Figure 1**b). Trials with >2 consecutive NoGo outcomes were taken to indicate disengagement and were discarded.

**Figure 1.**
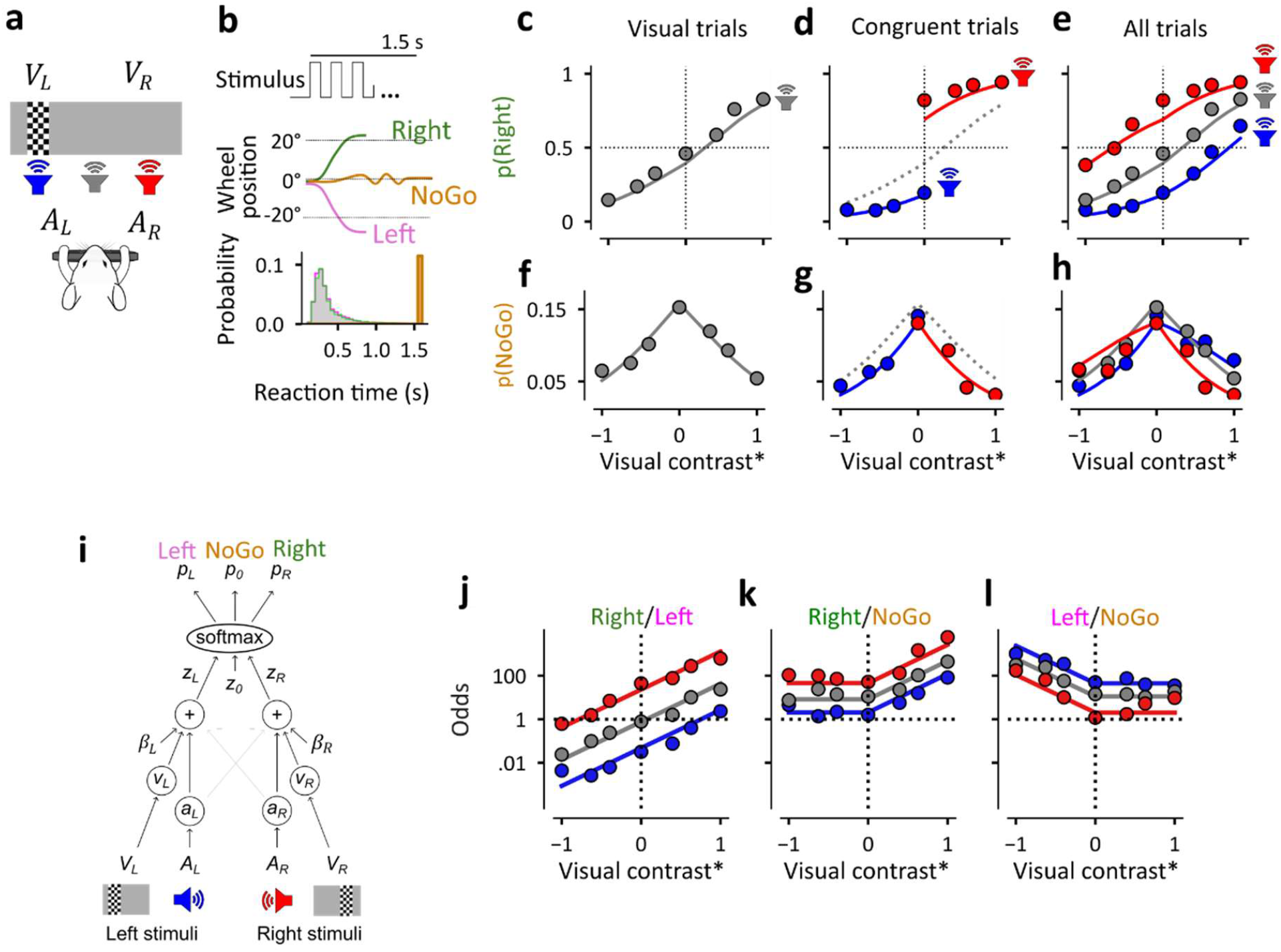
Mouse audiovisual choices obey 3-option logistic classification. **(a)** Task schematic. Visual and/or auditory stimuli are presented on the left or the right (*V*_*L*_, *V*_*R*_, *A*_*L*_, *A*_*R*_), unimodally, coherently, or in conflict. Mice turn a wheel to indicate their choices. **(b)** Top: Cartoon of the temporal task structure. Auditory and visual stimuli are pulsed coherently at 8 Hz until mice make a choice (≥20° wheel turn) or until the end of the 1.5 s response window. Middle: Wheel turn time trajectories. Bottom: Reaction time distributions (N=18 mice, subsampled for equal trial numbers (829 trials) per mouse). **(c)** Probability of rightward choices on visual trials at varying contrasts. Contrast is normalized (relative to the maximum contrast, which is 40% in these experiments) and raised by an exponent *γ* = 0.67, and is negative for stimuli on the left, positive for stimuli on the right, with auditory cue in the center (averaged over the same mice as in b). Curve is the fit of the model in **i. (d)** Same as **(c)**, for congruent trials, with auditory cues on the right (red) or left (blue), showing performance improvement relative to visual trials (dashed). **(e)** Performance on all trial types. **(f-h)** Fraction of NoGo choices in the same conditions as in **c-e**, showing dependence on trial difficulty. **(i)** The 3-option logistic classifier. Decision variables (*z*_*L*_, *z*_*R*_) depend on the weighted sum of sensory inputs (weights: *v*_*L*_, *v*_*R*_, *a*_*L*_, *a*_*R*_) and action pressure (*β*_*L*_, *β*_*R*_). Left, Right or NoGo choices are determined via softmax. **(j)** Odds of Right vs Left choices (in logarithmic scale) as a function of visual contrast (x-axis, rescaled by *γ*) and auditory location (color), showing fits of the model (lines). **(k-l)** Same, for the odds of Right vs NoGo, and Left vs NoGo.

Mice integrated the two cues, performing better in multisensory coherent trials than in unisensory trials (**Figure 1**c,d). Specifically, as seen in other multisensory tasks ^96–101^ the main effect of adding an auditory cue on the left or the right was to shift the visual psychometric curves to the left or the right ^14^ (**Figure 1**e).

### Mouse audiovisual choices obey 3-option logistic classification

In addition to Left and Right choices, mice made stimulus-dependent, unrewarded NoGo choices. Even after excluding consecutive NoGo trials, which may be associated with disengagement, we observed NoGo outcomes on 9.6 ± 1.2% of trials across mice (mean ± s.e., N = 18 mice across 491 sessions). These NoGo outcomes are unrewarded, but may be rational, because making actions is energetically costly. Indeed, they appeared to be non-random. First, they were unlikely to reflect late reaction times, because they were triggered after 1.5 s of inaction, and Left/Right choices occurred much earlier (303 ± 75 ms, median ± m.a.d., **Figure 1**b).

Second, they were unlikely to reflect missed trials, because every trial included a readily audible sound (to the left, middle, or right). Third, they depended on the stimuli, being more likely in difficult vs. easy trials: 4% higher in lower vs. top contrast in visual trials (p= 4.5*10^−6^, N = 18 mice, paired t-test, **Figure 1**f), 2% higher in visual vs. congruent audiovisual trials (p = 0.001, **Figure 1**g), and 3% higher in conflict vs. congruent audiovisual trials (p = 0.002, **Figure 1**h). The choice to withhold an action (NoGo) must have thus occurred after processing the stimuli. The probability of the three choices – Left, Right, and NoGo – obeyed a simple logistic model. To obtain the probabilities *p*(*L*), *p*(*R*), *p*(0) of Left, Right, and NoGo choices, we applied a 3-option logistic classification model ^2,10,15,16,102^ (**Figure 1**i), where

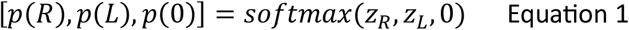

and *z*_*R*_ and *z*_*L*_ are two decision variables supporting Right and Left choices relative to a NoGo choice (whose decision variable is given the nominal value of zero). We explored multiple versions of the model (**Supplementary Figure 1**) and in the most successful version (**Figure 1**i) the decision variables have a simple linear dependence on the task variables:

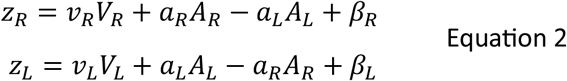

Here, *V*_*L*_ and *V*_*R*_ indicate the strength of the left or right visual stimulus (both zero when the visual stimulus was absent), and *A*_*L*_ and *A*_*R*_ indicate the left or right auditory stimulus (both zero when the auditory stimulus was in the middle). The parameters measure sensitivity for left and right visual stimuli (*v*_*L*_, *v*_*R*_), sensitivity for left and right auditory stimuli (*a*_*L*_, *a*_*R*_), and two stimulus-independent action pressures, for Left and Right actions (*β*_*L*_, *β*_*R*_).

With only 6 parameters, this model successfully described the probability of the three choices in 21 stimulus conditions. As expected ^14^ the model provided good fits to the probability of Right vs. Left choices (**Figure 1**e). In addition, it captured the prevalence of NoGo choices (**Figure 1**h). To inspect its performance, we plotted the log odds of all choice-pairs, as in this representation the model predictions become simple lines ^2,14^ (**Figure 1**j-l). For Left vs. Right choices, the model captured the linear effect of the visual stimulus (straight lines with slopes *v*_*L*_ and *v*_*R*_), and of the auditory stimulus (vertical shifts by *a*_*L*_ and *a*_*R*_) (**Figure 1**j). In addition, the model captured that the odds of Right choices relative to NoGo choices were independent of right visual stimuli (**Figure 1**k, flat line on the left), and vice versa (**Figure 1**l, flat line on the right). Auditory stimuli, instead, affected both the odds of Left and Right choices, relative to NoGo choices, and did so in opposite ways (**Figure 1**k and **Figure 1**l).

The model thus parsimoniously accounts for the data, while allowing us to estimate two separate pressures for Left and Right actions relative to inaction. By contrast, a previous 2-class model ^14^ applied to similar data ignored the NoGo choices and did not distinguish the two action pressures.

### Different SC neurons encode auditory and visual signals

To establish the potential role of the superior colliculus (SC) in this audiovisual task, we recorded from SC neurons with chronic^103^ Neuropixels probes^104,105^ during behavior and passive stimulus replay (**Figure 2**a, **Supplementary Figure 2**). To establish the location of each probe shank relative to the SC visual map, we mapped visual receptive fields using sparse noise stimuli (**Figure 2**b). As expected, the center of the screen mapped to anterior SC and the task visual cues were represented more posteriorly (**Supplementary Figure 2**b,d).

**Figure 2.**
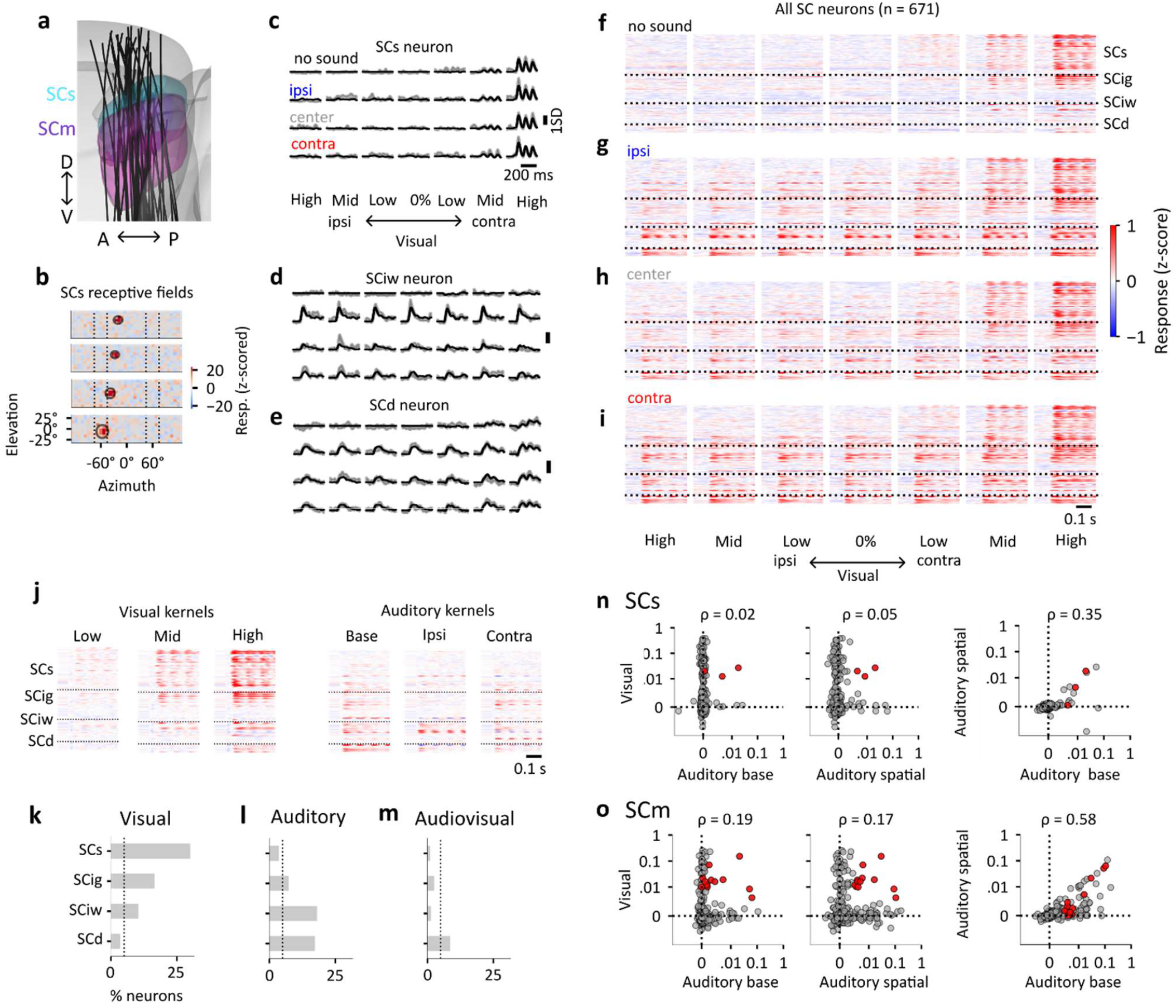
Different SC neurons encode auditory and visual signals. **(a)** Reconstructed probe trajectories registered to the Allen Atlas (46 shanks from 14 probes in N=9 mice); (**b)** Visual receptive fields (RFs) of four neurons in superficial SC (SCs) recorded in separate shanks in an example insertion, showing z-scored responses to white squares and fits of a 2D Gaussian. The stimulus locations are plotted for reference (dashed lines). **(c)** Responses of an example neuron in the superficial layer (SCs), plotted as mean ± s.e. (shaded) overlayed with model predictions (solid). Columns correspond to different visual contrasts (ipsi to contra) and rows to auditory stimulus presence and location (absent, ipsi, center, right). (**d**-**e**) Same, for two more example neurons in intermediate (SCiw) and deep layers (SCd). **(f)** Passive visual responses of all SC neurons (n=671) sorted by depth, showing borders (dashed lines) between layers. Each box shows responses at various contrast conditions (from ipsi to contra, as in c-e). No sound was played during these trials. **(g-i)** Same, when the auditory stimulus was played ipsilaterally to the neuron (g), in the center (h) and contralaterally (i). **(j)** Sensory response kernels for the same neurons, showing the 3 visual kernels (for 3 contralateral contrasts), the auditory base kernel (not directional), and the 2 directional auditory kernels (ipsi and contra). **(k-l)** Proportion of neurons with significant visual (k) and auditory (l) kernels in each SC layer. **(m)** Same, for neurons where both auditory and visual kernels are significant (audiovisual neurons). **(n)** Pairwise comparisons of variance explained in SCs: visual vs auditory spatial; visual vs auditory onset and auditory onset vs auditory spatial. Red dots: multisensory neurons **(o)** Same, for SCm (which consists of SCig, SCiw, and SCd).

We first examined the responses to task stimuli during passive replay, and as expected we found them to vary across layers. We presented a battery of auditory and visual cues that included those used in the task (setting their duration to 425 ms, which includes 4 stimulus pulses), and we recorded 671 well-isolated neurons that also remained stable during task performance (13 sessions from 9 mice, avoiding duplicate recordings by selecting one session per mouse from each brain region; **Supplementary Figure 2**c). Their responses were often pulse-locked to the 8 Hz stimuli, especially in the presence of high contrast contralateral visual stimuli (e.g., **Figure 2**c,e). Some neurons responded to sound (e.g., **Figure 2**d,e), occasionally with a preference for sound location (e.g., **Figure 2**d). As expected, these preferences were strongly depth-dependent, with responses to contralateral visual stimuli mostly in the superficial layer (SCs) and responses to sounds in SCm, which includes the intermediate granular, intermediate white, and deep layers (SCig, SCiw, and SCd, **Figure 2**f-i). To quantify these responses, we fitted each neuron’s activity with a linear model based on visual and auditory kernels. The model included three contralateral visual kernels *V*_1_, *V*_2_, *V*_3_ (for low, middle, and high contrast), and three auditory kernels *A*_*b*_, *A*_-1_, *A*_1_ (base, ipsilateral, and contralateral) (**Supplementary Figure 3**b). According to the model, the sensory responses of a neuron at time *t* (relative to stimulus onset) in trial *j* are given by

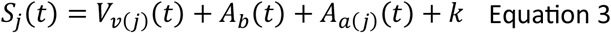

where *v*(*j*) indexes the contralateral contrast presented in trial *j* (low, middle, or high), and *a*(*j*) indexes the auditory position (contralateral or ipsilateral), and *k* is a constant. The first term applies when the visual stimulus was contralateral. The second term applies when any auditory stimulus was present. The third term applies when the auditory stimulus was present and not in the center.

The model provided good fits (e.g. **Figure 2**c-e) with visual kernels strongest in SCs and auditory kernels more likely in deeper layers (**Figure 2**j, **Supplementary Figure 3**c,d). We measured each kernel’s unique contribution by calculating the cross-validated variance explained after subtracting all other kernels’ predictions. Using a threshold of >0.5% variance explained, we found that 20% of SC neurons were visual and 10% were auditory (average across mice). Visual neurons concentrated in superficial layers (SCs: 31%, SCm: 15%, estimated across mice with linear mixed effects model direction of difference tested with one-sided Wald test, N=9 mice, p=0.037), **Figure 2**k, **Supplementary Figure 3**g) and were typically driven (87%) rather than suppressed (13%) by contrast. Auditory neurons were spatially segregated, appearing predominantly in SCm (SCs: 4%, SCm: 14%, N=9 mice, p = 0.020, **Figure 2**l, **Supplementary Figure 3**h), and typically (66%) preferred contralateral sounds (the rest preferred ipsilateral sounds or were untuned for sound location (**Supplementary Figure 3**h).

In contrast with many reports^21–31^ emphasizing audiovisual integration in SC, most of the responses were strikingly unimodal: only 3% of all SC neurons (SCs: 1.3%, SCm: 3.9%) were both auditory and visual (**Figure 2**m). To compare auditory and visual contributions to the activity of individual neurons, we compared the variance explained by the three visual kernels to the variance explained by the auditory base kernel or by the two auditory spatial kernels (ipsi and contra). For most neurons, the variance in the responses was explained by the visual kernels alone or by the auditory kernels alone (**Figure 2**n,o). The variance explained by the two was not correlated in SCs (ρ = 0.02 and 0.05 for the base and spatial auditory kernels, p = 0.71 and p = 0.38, hierarchical Spearman’s correlation from mixed effects rank regression) and was significantly but weakly correlated in SCm (ρ = 0.19 and 0.17, p = 0.0001, and p = 0.0006). In contrast, the variance explained by the two types of auditory kernel (base and spatial) was strongly correlated (ρ = 0.35 in SCs; ρ = 0.58 in SCm, p = 9.1*10^−10^ and p = 6.5*10^−45^). Overall, these results indicate that visual and auditory signals remain most often separate in the SC, and neural responses to audiovisual stimuli are largely dominated by one sense, with weak modulation across the senses.

### Different SC neurons encode sensory and task signals

We next examined the responses of the same neurons during the task and quantified the correlates of task engagement and upcoming actions. Engagement and actions can modulate the activity of neurons all over the brain (including SC) ^106^. To capture these effects in our SC population, we extended the linear model we had already fitted to the passive responses. In addition to the six sensory kernels, which were the same as in the passive condition (**Figure 3**a) the extended model included two stimulus-independent kernels: an engagement kernel *E* (**Figure 3**b) and two action kernels *C*_-1_, *C*_1_ for ipsilateral and contralateral actions (**Figure 3**c).

**Figure 3.**
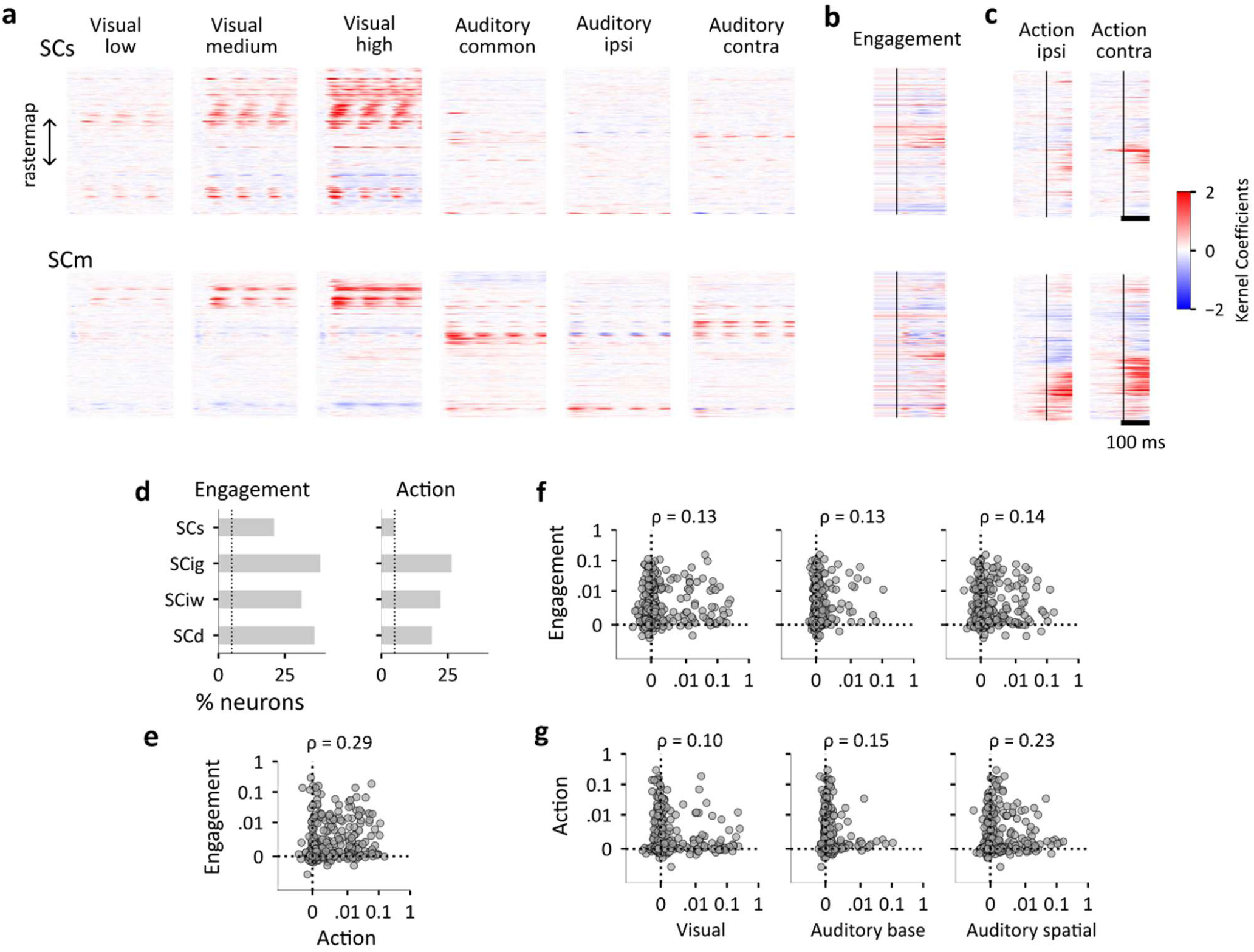
Different SC neurons encode sensory and task signals. **(a)** Sensory response kernels of SCs (top, n=276) and SCm (bottom, n=375) neurons. These are the same kernels as Figure 2j, but here neurons are sorted by Rastermap^107^ rather than by depth, to better visualize the kernels. **(b)** Engagement kernels for the same neurons **(c)** Action ipsi and contra kernels for the same neurons. Action kernels are aligned to wheel-movement onset (solid line), and engagement and stimulus kernels are aligned to stimulus onset. **(d)** Fraction of neurons with significant engagement and action kernels per layer. **(e)** Comparison of variance explained by engagement vs. action kernels in SCm neurons. **(f)** Variance explained by sensory vs. task kernels in SCm neurons. **(g)** Variance explained by sensory vs. action kernels in SCm neurons.

As with the passive case, the extended model is linear: each kernel provides convolution and the resulting terms are added to each other. In the model, the task responses of each neuron at time *t* in trial *j* are given by

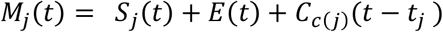

where *S*_*j*_ is the sensory response of the neuron to the stimuli presented in trial *j* (measured in the passive stimulus presentations, Equation 3), *E* is the engagement kernel (which depends on time but is the same across all trials in a task session), and *c*(*j*) = ±1 indicates the action made in trial *j*, and *t*_*j*_ is the time of that action. If no action was taken (NoGo), the third term does not apply.

The engagement kernel captures overall differences in activity between task and passive presentation, which could occur before and after stimulus presentation. It tended to be constant across time (explaining 0.7% of variance, while the time-dependent modulation explained 0.1% variance on average across neurons) and was slightly more positive (in 65% of SCs engagement neurons and 60% of SCm engagement neurons, and negative in the rest, **Figure 3**b).

The two action kernels were also usually both positive (79% of ipsi kernels, and 86% of contra kernels), and many neurons contained both, as they were positively correlated (ρ = 0.36, p = 1.9*10^−24^) (**Figure 3**c). The contralateral action kernel on average was slightly stronger than the ipsilateral one across mice^106^ (contra – ipsi amplitude was > 0 in 61% of action neurons, one sample t-test on the kernel amplitude difference for each action neuron, N=9 mice, p=0.02).

While signals related to engagement occurred in all layers, signals related to actions were specific to SCm. In 58% of the SC neurons, the extended model explained responses better than a constant mean firing rate (cross-validated R^2^>10^−3^), indicating that these neurons were modulated by stimuli, engagement, or actions. Using the variance explained score on the residuals (>0.5% threshold), we found that 33% of SC neurons were modulated by engagement, and 16% by actions. Engagement modulation occurred in all layers, but more so in SCm (SCs: 21%, SCm: 37%, N=9 mice, p = 0.007 **Figure 3**d, **Supplementary Figure 3**i) Action neurons were even more strongly localized to SCm (SCs: 6%, SCm:20%, N=9 mice, p = 0.002, **Figure 3**d, **Supplementary Figure 3**j). In SCm, 12% of neurons were modulated both by engagement and actions, and engagement and action modulation was correlated (ρ = 0.29, p = 2.3*10^−9^, **Figure 3**e).

The modulation of SCm neurons by engagement and action was independent of sensory tuning. Though 33% of neurons were modulated by stimuli, only 15% were modulated by both stimuli and engagement **Figure 3**f), and only 7% by both stimuli and actions (**Figure 3**g). These numbers resemble those expected by chance (12% and 7%). Indeed, the correlations between variance explained by engagement, action, and sensory kernels were significant but small (all ρ<0.25 and p>2.1*10^−6^), with most variance explained by action kernels or by sensory kernels but rarely by both (**Figure 3**f,g).

Taken together, our observations during passive presentation (**Figure 2**) and during task performance (**Figure 3**) argue against a prevalence in mouse SC of integrative computations such as audiovisual integration or sensorimotor transformation.

### SCm and MOs activity helps predict trial-by-trial choices

To assess the trial-by-trial relationship between neural activity and choices, we turned to a decoding approach. We extended the logistic model so that its inputs include not only the sensory stimuli, but also neural activity in a given region (**Figure 4**a). In the extended model, the decision variables are given by

**Figure 4.**
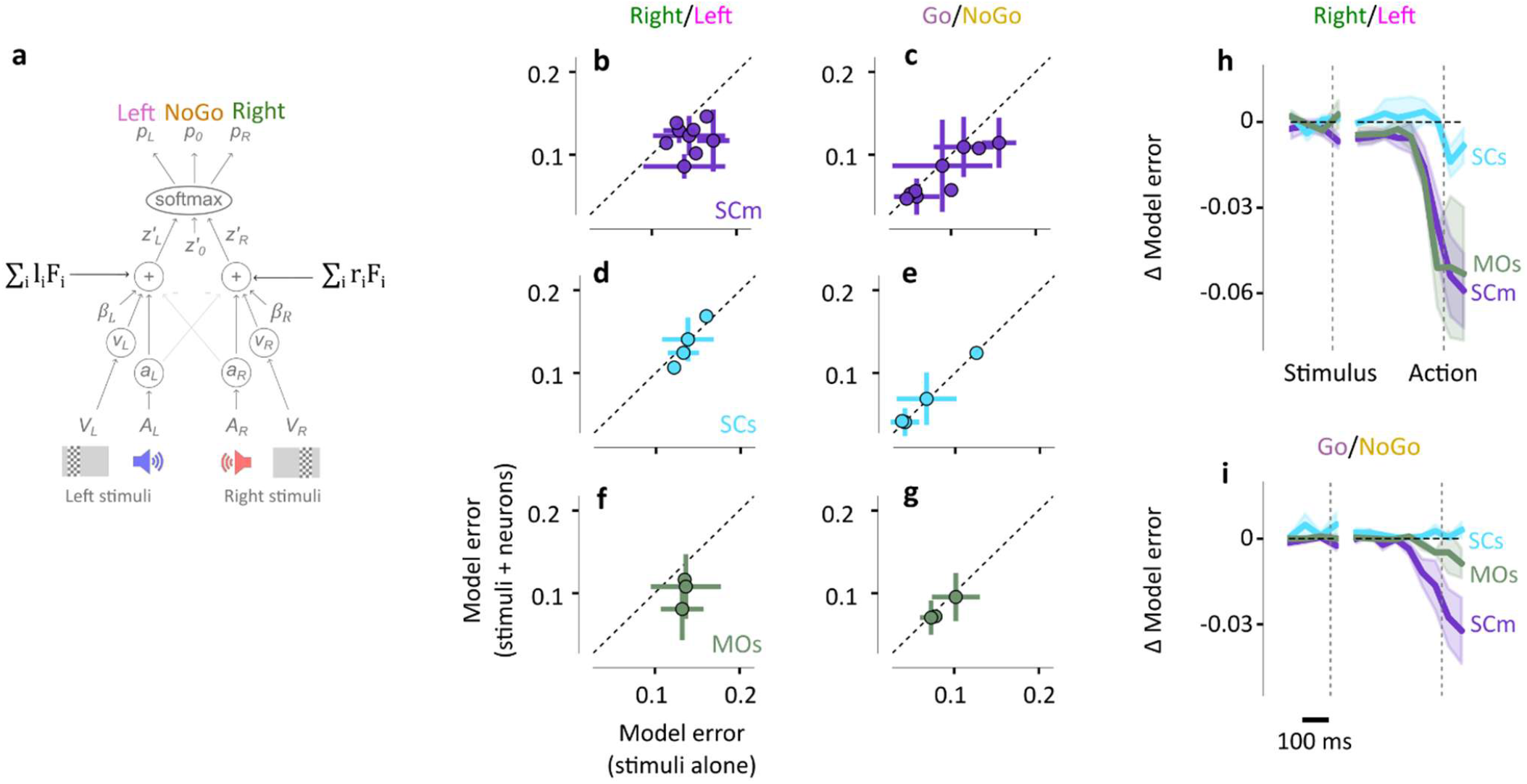
SCm and MOs activity helps predict trial-by-trial choices. **(a)** Schematic of the 3-option neural decoder, showing the stimulus-based model (*gray*) and the additional neural predictors (*blue*). **(b)** Average error for the stimulus-only model (x-axis) versus the model including both stimuli and SCm activity (y-axis), for Right vs Left choices. Points represent mice, averaged across sessions (N=9 mice, 26 sessions). Error bars indicate ±1 S.D. across sessions. Values below the identity line indicate improved prediction when including neural activity. **(c)** Same, for Go vs NoGo choices. **(d-e)** Same as b-c but using SCs neural activity (N=4 mice, 6 sessions). **(f-g)** Same as b-c but using MOs neural activity (N=3 mice, 19 sessions). **(h)** Reduction in model error when predicting Left/Right choices when using neural activity from SCm (*purple*), MOs (*green*), or SCs (*cyan*), as a function of time. Dashed lines indicate times of stimulus and action. **(i)** Same, for Go/NoGo choices. Curves indicate averages across mice; shaded areas denote ±1 s.e. across mice. Negative values indicate improved predictions.

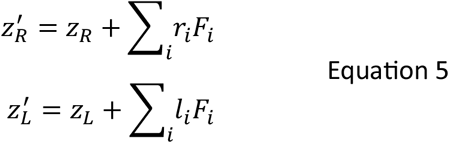

where *z*_*L*_ and *z*_*R*_ are the decision variables defined earlier based on stimulus attributes (Equation 2), the sum is over neurons *i* in a given region, *F*_*i*_ are their firing rates calculated in a 200 ms window prior to action, and *l*_*i*_ and *r*_*i*_ are weights scaling each neuron’s contribution to the two decision variables. Reasoning that neurons encoding action or engagement are those most likely to contribute to choice improvement, we restricted our decoder to only include such neurons (relaxing this assumption did not change the results). We then evaluated model performance for two choice components: the decision to act (Go vs NoGo), and the direction of that action (Left vs Right).

Unlike the original model, this extended model can predict different choices in trials that have the same sensory stimuli. We thus tested whether including the neural activity improved the predictions on a trial-by-trial basis. To evaluate the performance of the models (with and without neural activity) we calculated their average error (Brier score^108^): the mean squared difference between predicted probability and actual outcome expressed as 0 or 1. Similar results were obtained with a measure that characterizes both accuracy and confidence (the functional margin, **Supplementary Figure 5**).

Population activity in SCm, but not SCs, provided robust trial-by-trial predictions of the upcoming choices. Adding SCm activity to the stimulus-based model consistently increased the accuracy of predictions (reduced average errors) for the direction of the action (Right vs Left, p=0.0013, linear mixed effects model with one-sided Wald z-test, **Figure 4**b) and for the action itself (Go vs NoGo, p = 0.0065, **Figure 4**c). By contrast, SCs activity did not improve the predictions (p = 0.289 for Right vs Left, p = 0.907 for Go vs NoGo, **Figure 4**d,e).

Choice decoding from SCm activity was as good as that seen in prefrontal area (MOs). We recorded from 498 neurons in MOs in 3 mice that performed the task and found neurons with both action and engagement kernels (**Supplementary Figure 4**). Incorporating MOs activity into the model improved both the confidence and accuracy of Left/Right predictions (p = 0.0001, **Figure 4**f, **Supplementary Figure 5**f). For Go/NoGo choices, however, MOs activity mostly improved model confidence, barely improving accuracy (p = 0.048, **Figure 4**g, **Supplementary Figure 5**g). The contributions by MOs and SCm can be most directly compared when overlaid as a function of time (**Figure 4**h,i). Before stimulus onset, neither region contained activity that predicts upcoming action. About 100 ms before action, both MOs and SCm activity improved Left/Right predictions with similar accuracy and timing; this improvement (reduction in average error) was not significantly different between the two regions (p=0.462, linear mixed effects model, with decoder using averaged activity 0-200 ms prior to choice, **Figure 4**h). Similarly, the improvement in predicting Go/NoGo choices did not statistically differ between SCm and MOs (p = 0.316, **Figure 4**i).

### Each side of SCm promotes contralateral action relative to inaction

These neural signals suggest that SCs is unlikely to play a role in determining the choices beyond a possible sensory role, whereas SCm could play a variety of roles, comparable to MOs. However, the presence of neural signals does not demonstrate a causal link to behavior.

To establish the causal role of the SCm in the task, we turned to optogenetic inactivation. We injected *hSyn-halorhodopsin* (eNpHR3.0) ^109^ into the intermediate and deep layers of the SC (unilaterally or bilaterally) and inserted one or two cannulae to target the top of the injection sites (10 mice, **Supplementary Figure 6**a). Inactivation trials (35%) were randomly interleaved with control trials. Recordings performed during inactivations in control mice confirmed a decrease in neural activity near the cannula site (**Supplementary Figure 7**).

Unilateral SCm inactivation markedly decreased contralateral choices in favor of inaction, regardless of sensory stimuli. For example, inactivation of left SC markedly decreased the fraction of Right choices (**Figure 5**a). The missing Right choices, moreover, were replaced mostly by NoGo choices (**Figure 5**b), and rarely by Left choices (**Figure 5**c). Crucially, these effects were present regardless of the position of auditory and visual stimuli, and regardless of visual contrast. Symmetric effects were seen when inactivating right SC (**Figure 5**d-f), so in the subsequent analyses we pooled them, denoting the effects as contralateral (c) or ipsilateral (i) to the inactivated site. For instance, the parameter *β*_*C*_ refers to *β*_*R*_ when inactivating left SC, and to *β*_*L*_ when inactivating right SC.

**Figure 5.**
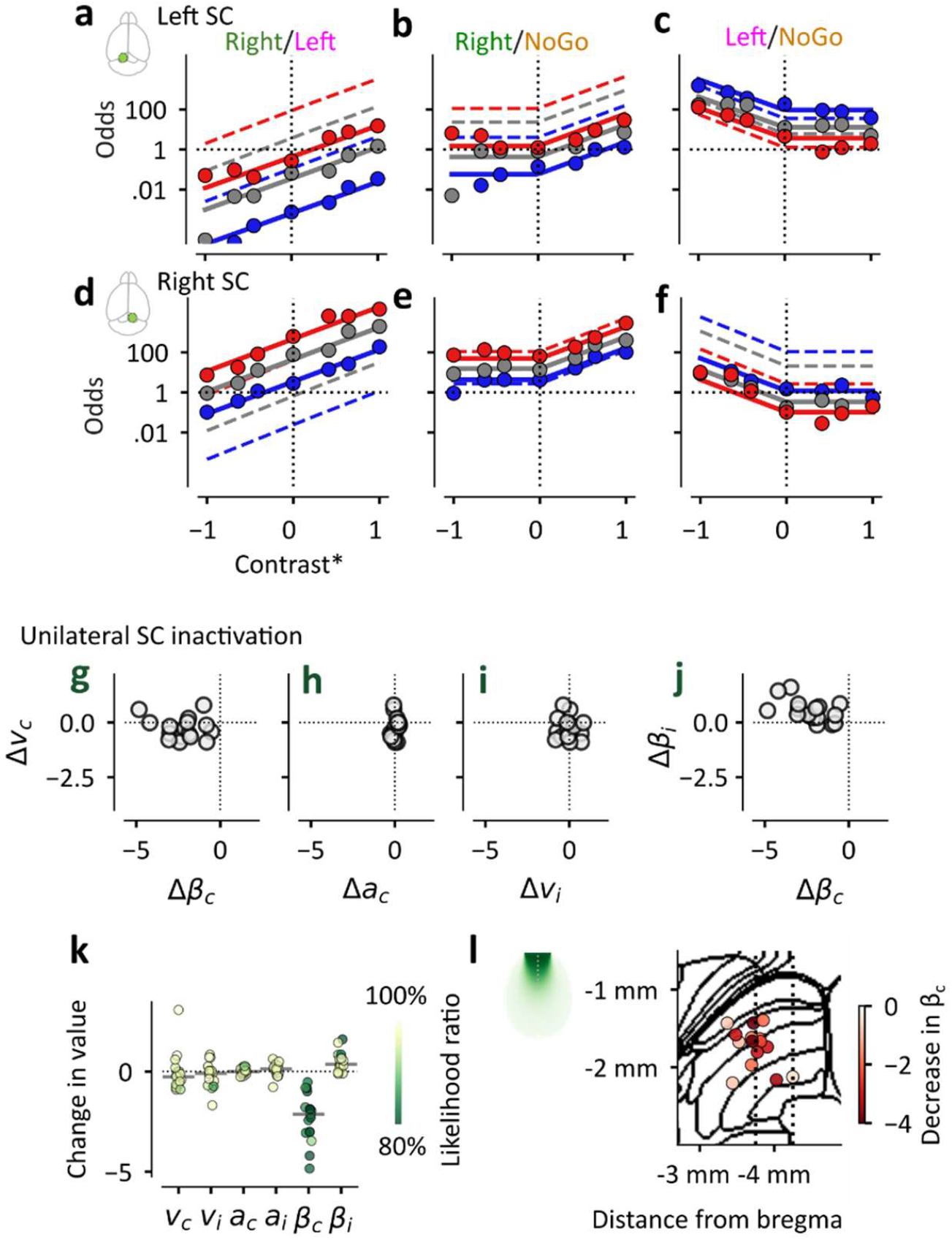
Each side of SCm promotes contralateral action relative to inaction. **(a-c)** Effects of inactivating the left SC on the odds of Right vs Left (a) Right vs NoGo and (b) on the Left vs NoGo (c) choices. N=8 mice, averaging 1,316 trials/mouse. **(d-f**) Same as (a-c) for right SC inactivation. N=9 mice, average of 4,923 trials/mouse; 7 mice contributed to both left and right datasets via bilateral implants. (**d-g**) Effects of unilateral SC inactivation on model parameters: contralateral action pressure *β*_*C*_ and contralateral visual sensitivity *v*_*C*_ (d), contralateral auditory sensitivity *a*_*C*_ (e), ipsilateral visual sensitivity *v*_*i*_ (f), and ipsilateral action pressure *β*_*i*_ (g). Each dot is a hemisphere (17 hemispheres from 10 mice). One hemisphere is omitted because its values for the *v*_*C*_ parameter is >2.5 (see panel k). **(k)** Changes in model parameters induced by unilateral SC inactivation. Same data as in d-g, colored by the confidence in the parameter contribution, assessed via likelihood ratio: the loss in log-likelihood relative to the extended model when that parameter is dropped. **(l**) Estimated spatial extent of the inactivation effect around the cannula tip (left) and diagram of the SC in the sagittal plane showing cannula positions (right). The color indicates the change in contralateral action pressure *β*_*R*_.

The 3-option logistic model (Equation 1-2) captured these effects mostly through a decrease in the contralateral action pressure. Inactivation of one side of SC sharply reduced the contralateral action pressure term *β*_*C*_ while not affecting contralateral visual sensitivity *v*_*C*_ (**Figure 5**g), contralateral auditory sensitivity *a*_*C*_ (**Figure 5**h), or ipsilateral visual sensitivity *v*_*i*_ (**Figure 5**i). The strong reduction in contralateral action pressure *β*_*C*_ was accompanied by a mild increase in ipsilateral action pressure *β*_*i*_ (**Figure 5**j), resulting in a slight increase in overall ipsilateral choices relative to NoGo choices (visible e.g. in **Figure 5**c).

To assess the effect of inactivations on the parameters of the model, we measured the likelihood ratio of model prediction obtained while allowing that parameter to vary vs. while holding it fixed (**Figure 5**k). Unilateral SC inactivation significantly reduced the contralateral action pressure term, which we term *β*_*C*_ (p = 6.5*10^−7^, paired t-test between control and inactivation parameters, N = 17 hemispheres (from 10 mice)

**Figure 5**k). This decrease did not visibly depend on the exact position of the optic fiber (**Figure 5**l, **Supplementary Figure 6**a,b). It was accompanied by the aforementioned small increase in ipsilateral action pressure *β*_*i*_ (p = 0.0004, **Figure 5**k). There was, however, no effect on visual weights (p = 0.75 for *v*_*C*_ and p = 0.35 for *v*_*i*_) or on auditory weights (p = 0.61 for *a*_*C*_ and p = 0.47 for *a*_*i*_): keeping these weights constant across inactivation conditions did not decrease fit quality (**Figure 5**k).

In this task, therefore, the SC does not contribute to audiovisual localization; rather, each side of SC provides a stimulus-independent pressure that mostly promotes the contralateral choice and slightly suppresses the ipsilateral choice. As explained by our 3-option model, this pressure is only revealed during unilateral inactivation, because it is otherwise balanced by an equal pressure from the other side. In normal conditions, the two equal pressures equate the chances of Left and Right choices and reduce those of NoGo choices.

### The two sides of SCm provide additive contributions

This understanding of the role of SC makes three simple predictions for the effects of bilateral SC inactivation. First, analogous to the Sprague effect ^90,110,111^, inactivating both sides of SC equally should restore the balance of Left and Right choices. Second, it should do so largely by making both choices less likely relative to NoGo choices. Third, the effects of bilateral inactivation should be additive, i.e. equal to the sum of unilateral inactivations.

To test the first two predictions, we inactivated the two sides of SC with approximately equal intensities and found that both predictions were correct. First, inactivating both sides of SC restored the balance between Left and Right choices (**Figure 6**a). Second, this balance occurred because both Right and Left choices became less likely relative to NoGo choices (**Figure 6**b,c).

**Figure 6.**
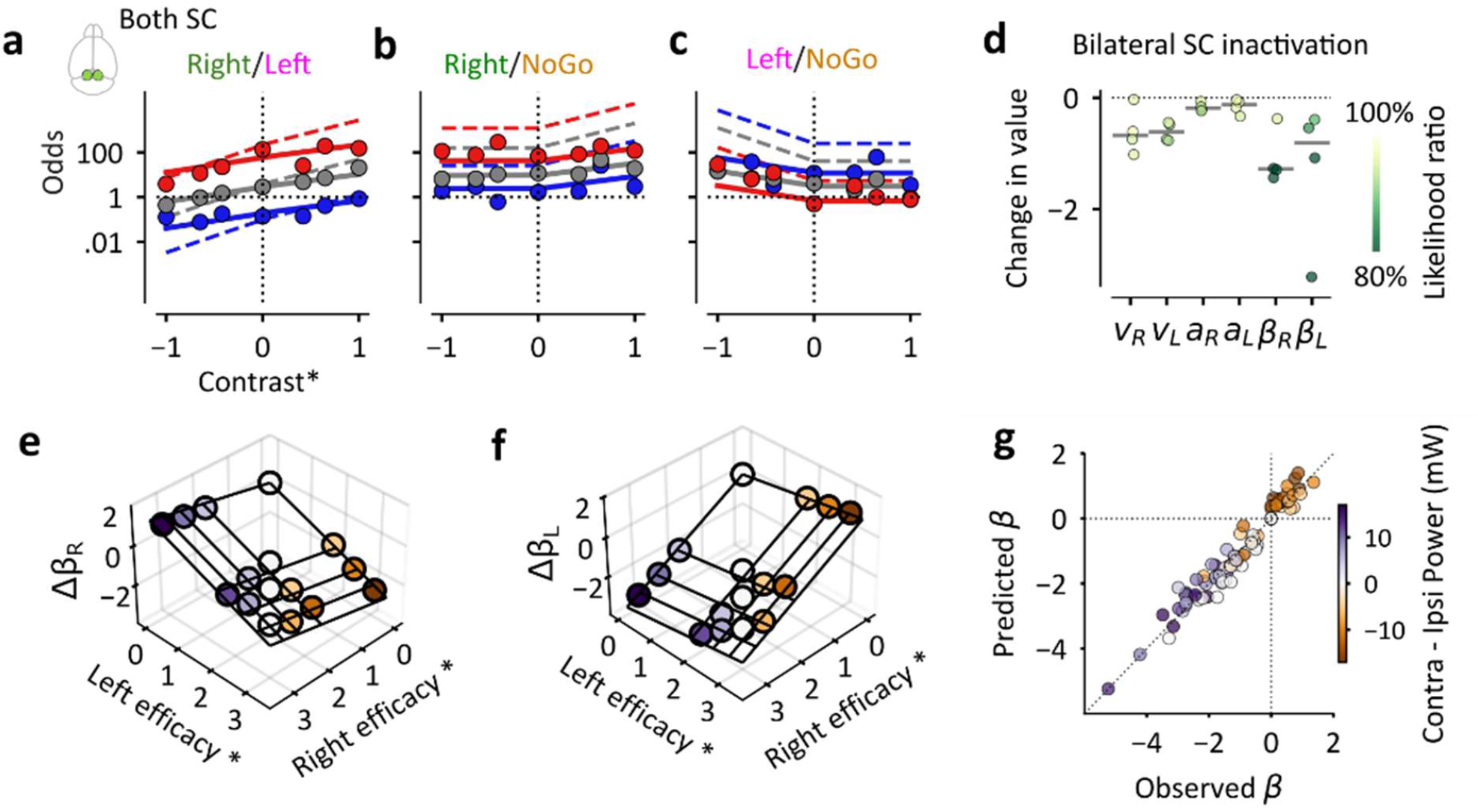
The two sides of SCm provide additive contributions. **(a-c**) Effects of bilateral SC inactivation on choice Odds at high powers (10-17 mW/mm^2^, equal across hemispheres) (4 mice, 4,384 trials/mouse trials subsampled to be balanced across mice). (**d**) Changes in model parameters induced by bilateral SC inactivation, colored by the confidence in the parameter contribution, formatted similarly to Figure 5k. (**e**). Changes the right action pressure (Δ*β*_*R*_) for an example mouse, as a function of inactivation efficacy in the left hemisphere (x-axis) and the right hemisphere (y-axis). Dots are empirical Δ*β*_*R*_estimates, and the solid line is the fit obtained by additive power model in Equation 6. Each dot is colored by the difference in inactivation powers (Right – left). **(f)** Same, for the left action pressure (Δ*β*_*L*_). **(g)** Observed vs. predicted action pressure parameters (Δ*β*_*L*_ and Δ*β*_*R*_) across all power combinations in out-of-sample test data (4 mice). Colors indicate the difference in contra – ipsi inactivation power. To evaluate predictive accuracy, trials were split into two independent halves and the empirical Δ*β* parameter was obtained on each. Including ipsi parameters resulting in significantly better fits (p=0.009 paired t-test on the R^2^ score of the test set).

The 3-option model captured these effects largely via a reduction in action pressure terms *β*_*L*_ and *β*_*R*_ (**Figure 6**d). This reduction was significant (p = 0.01, repeated measures ANOVA, N = 4 mice) and not different between the two sides (p = 0.11). This effect was highly robust: imposing no change in either action pressure greatly decreased the likelihood of the model fits (average decrease in likelihood ratio across both *β* parameters was 27% **Figure 6**d). In addition, bilateral inactivation slightly decreased the visual and auditory sensitivity terms (p = 0.009 and p = 0.042 **Figure 6**d). However, these decreases did not seem as robust as the decrease in action pressure terms *β*_*L*_ and *β*_*R*_: imposing no change in visual sensitivity or auditory sensitivity barely affected the likelihood of the model predictions (average decrease in likelihood ratio was 2% and 1%).

To test for additivity, we performed bilateral inactivations while systematically varying the light intensity of the inactivation in each hemisphere. The action pressure for Right choices, *β*_*R*_, was markedly decreased by light in the left hemisphere, and mildly increased by light in the right hemisphere (**Figure 6**e). The symmetrical effects were observed in the action pressure for Left choices, *β*_*L*_ (**Figure 6**f). To formalize these effects, we modeled Δ*β*_*L*_ and Δ*β*_*R*_ with a simple linear model:

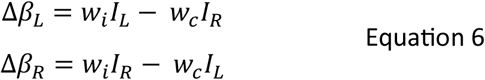

where *I*_*L*_ and *I*_*R*_ are the inactivation efficacies achieved in the left and right SC (laser power scaled by a constant for each hemisphere and raised to a shared exponent to capture power-nonlinearities, see Methods), and *w*_*C*_ and *w*_*i*_ indicate the effect of contralateral and ipsilateral light on each action pressure (contralateral suppression and ipsilateral facilitation). The model provided good fits to the data, both in an example mouse (**Figure 6**e-f) and across mice (**Figure 6**g). As we have seen with unilateral SC inactivations (**Figure 5**k), the estimated value for the effects of contralateral suppression was much larger than the ipsilateral facilitation (*w*_*C*_ ≫ *w*_*i*_).

### The SCm promotes competing temporal priorities

Unilateral SCm inactivations, but not bilateral ones, also impacted reaction times. Unilateral inactivation sped up ipsilateral choices by 68 ± 37 ms (median ± m.a.d., averaged across mice and stimulus conditions first, from N = 17 hemispheres from 10 mice, n = 128,262 trials, of which 21,359 were inactivation trials), a significant decrease (p = 1.5*10^−5^, Wilcoxon signed rank test against 0, N = 17). Conversely, it slowed down contralateral choices by 99 ± 52 ms (p = 4.6*10^−5^). These effects canceled out in the bilateral inactivations, which did not alter reaction times (p = 0.55, change in reaction times: 8 ± 46 ms, N = 4 mice, n = 35,283 trials, where 7,144 were inactivation trials).

Each side of SCm, therefore promotes contralateral action not only by making it more likely but also by making it faster. When the SC is intact, the opposing effects provided by the two sides cancel each other, making the two choices more equally likely and more equally timed.

To understand the SC’s effect on both choices and reaction times, we turned to the well-established drift-diffusion model ^12,112–114^ (DDM) and extended it to account for the varying prevalence of NoGo choices in our experiments. At each timepoint, a DDM^12,112–114^ performs a step of the logistic classification ^2^, with visual and auditory inputs additively contributing to the drifting variable together with a constant drift bias. NoGo outcomes stem from trials that reach neither boundary within a set time. To account for their large fraction in the data, we postulated a ‘lazy signal’ gain function that multiplies the evidence-driven drift rate and decreases over time (opposite to an ‘urgency signal’ ^115–117^). The model provided good fits suggesting that mice prioritized sensory evidence presented early in the trials ^118^, and choose inaction when the resulting judgement of reward odds was unfavorable (**Supplementary Figure 8**).

The extended DDM confirmed the causal role of SC in promoting competing actions both in prevalence and in timing. The modified DDM captured the effects of inactivation through two concurrent changes (**Supplementary Figure 9**) a change in drift bias (as in Ref. ^119^), which explained the changes in reaction time seen during unilateral inactivation, and a change in the decision boundary (as in Ref. ^45^), which explained the increase in Inaction choices. These results potentially reconcile earlier results obtained in primates ^30,79^ and again argue that contributions of the SC to these decisions is independent of sensory evidence.

### Visual and prefrontal cortex play sensory and integrative roles

To further interpret the results of SC inactivations, we compared them to unilateral inactivations in visual and prefrontal cortical areas. We analyzed the results of optogenetic inactivations of left visual cortex (VIS) or left prefrontal cortex (MOs), obtained by stimulating ChR2 in inhibitory PV cells ^14^, and fitted the results with the three-class logistic model. As shown previously ^14^, both inactivations decreased Right choices relative to Left choices (**Figure 7**a,d). The analysis of the NoGo trials, moreover, revealed that (as with unilateral SC inactivations) this impairment in Right choices was accompanied by an increase in NoGo choices (**Figure 7**b) rather than an increase in Left choices (**Figure 7**c). However, unlike in SC, inactivations of left VIS impaired Right choices only when the visual stimulus was on the right (**Figure 7**a,b), suggesting that the mouse had difficulty perceiving contralateral visual stimuli and acted as if they were absent. By contrast, inactivations of left MOs impaired Right choices in all trial types (**Figure 7**d,e). In addition, they slightly decreased Left choices relative to NoGo choices when visual stimuli were on the left (**Figure 7**f).

**Figure 7.**
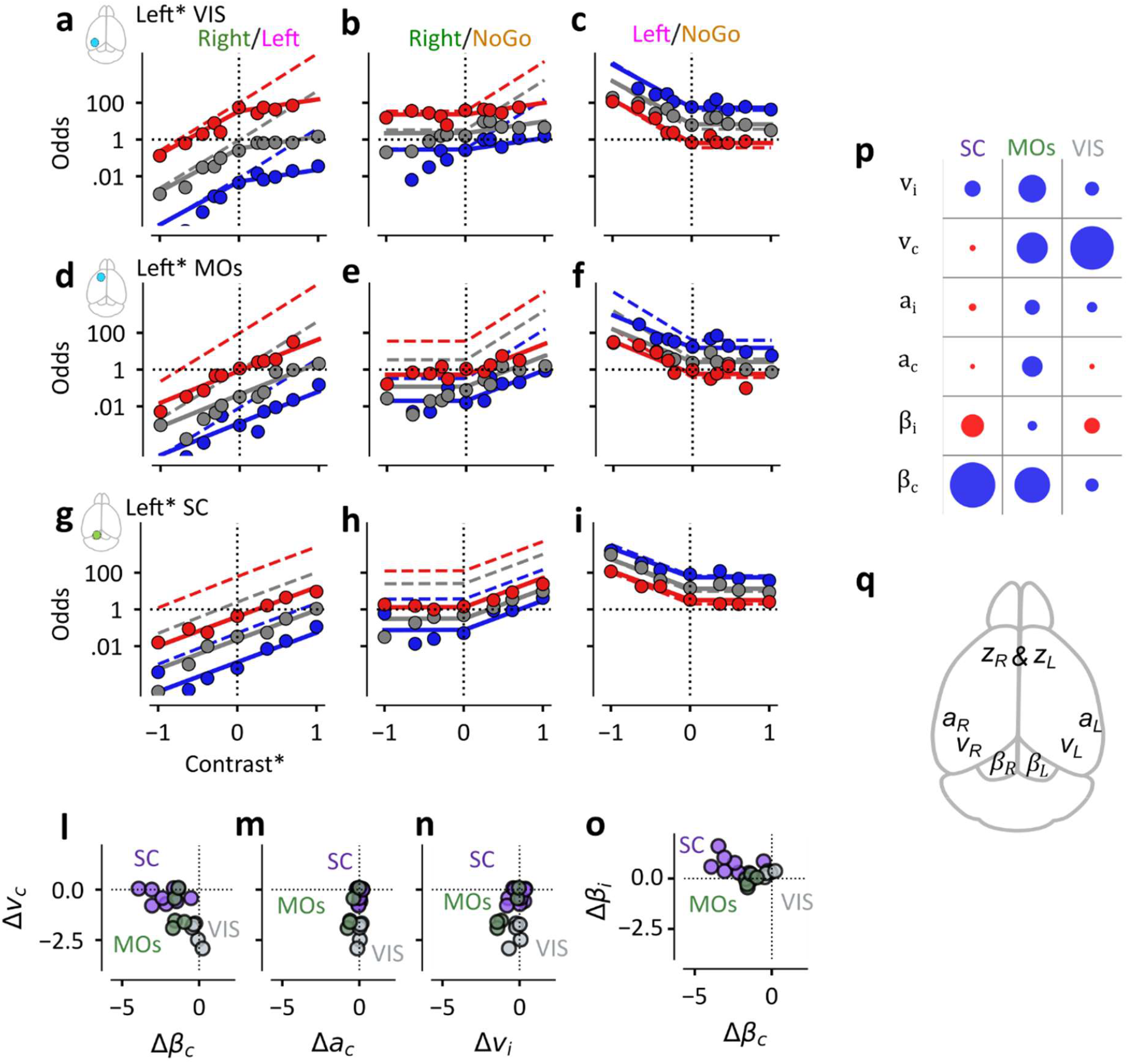
Visual and prefrontal cortex play sensory and integrative roles. **(a)** Odds of choosing Right over Left after inactivation of one side (nominally the left side) of visual cortex (VIS) via PV-ChR2.0, N=5, 2,572 trials/mouse, showing model fits on inactivation trials (solid lines) and on control trials (dashed lines). Data from left and right inactivations are combined and plotted as appropriate for left inactivations. **(b-c)** Same, for the odds of Right over NoGo (b) and Left over NoGo (c). **(d-f)** Same as (a-c), after unilateral inactivation of prefrontal cortex (MOs), N=5 mice (same mice as in (a-c), 2604 trials/mouse; **(g-i)** Same, after unilateral inactivation of SC. N=10 mice, 1316 trials/mouse. **(l)** Changes in contralateral visual sensitivity *v*_*C*_ vs. action pressure *β*_*C*_, after unilateral inactivation MOs (green) and VIS (gray), and SC (purple) across mice; When data from unilateral inactivation of both hemispheres was available, we took the mean across the two numbers. **(m-n)** Same but plotted against changes in contralateral auditory sensitivity *a*_*C*_ (h) and in ipsilateral visual sensitivity *v*_*i*_ **(o)** Same, comparing ipsi-vs contralateral action pressure (*β*_*i*_ vs. *β*_*C*_) **(p)** Summary of the average effect of unilateral SC, MOs or VIS inactivation on visual, auditory and action pressure parameters. The area of each dot indicates the average effect across mice (blue for decreases, red for increases). **(q)** Our results suggest that primary sensory cortices are responsible for perceiving unisensory sensory evidence on the contralateral hemifield, the frontal cortex integrates all decision variables, and the SC promotes available actions.

These effects of unilateral inactivations in the cortex differ from those seen in SC (replotted in **Figure 7**g-i for convenience), supporting a purely visual role for visual cortex and an integrative role for prefrontal cortex. The effects were captured by the logistic model (**Figure 7**a-i), so once again we can describe them through the changes in the model’s parameters. Inactivation of VIS only decreased contralateral visual sensitivity *v*_*c*_ (−2.1 ± 0.2, mean ± s.e.), with minimal effects on contralateral action pressure *β*_*C*_ (−0.2 ± 0.1) and the other parameters (**Figure 7**l-n). By contrast, inactivation of MOs decreased not only contralateral visual sensitivity *v*_*R*_ (−1.1 ± 0.4) but also contralateral action pressure *β*_*C*_ (−1.4 ± 0.1, **Figure 7**l) and auditory sensitivity *a*_*R*_ (−0.5 ± 0.1, **Figure 7**m), as well as ipsilateral visual sensitivity *v*_*i*_ (−0.9 ± 0.3 **Figure 7**n). All these effects favor NoGo choices, especially for stimuli on the right.

By providing nonsensory action pressures, therefore, the SCm is in a unique position relative to visual and prefrontal areas. Unilateral inactivation of SCm decreased contralateral action pressure *β*_*C*_ by −2.3 ± 0.3 (mean ± s.e.), significantly more than inactivation of MOs and VIS (p= 0.001, one way ANOVA). Conversely, it left contralateral visual sensitivity *v*_*c*_ unaffected (an average increase of 0.0 ± 0.4), a very different effect from the strong decreases seen when inactivating VIS or MOs (p = 0.002, one way ANOVA). Finally, inactivation of SCm - but not MOs or VIS – slightly increased ipsilateral action pressure *β*_*i*_(SC: 0.6 ± 0.2, MOs: −0.1 ± 0.1, VIS: 0.3 ± 0.1, p=0.019, one way ANOVA, **Figure 7**o).

To ensure the differences between SC and MOs were not due to differences in the inactivation method — increased inhibition via activation of PV neurons with ChR2 in MOs vs. direct hyperpolarization with halorhodopsin in SC – in a separate cohort of mice we repeated the MOs inactivation using halorhodopsin. Even with the matched opsin, the effects of MOs inactivation remained strikingly different from those of SC inactivation (**Supplementary Figure 10**): it reduced visual (p=0.0006, repeated measures ANOVA, N=4 mice) and auditory sensitivity (p=0.02) but not contralateral action pressure (in fact, it slightly increased it, p = 0.03, paired t-test, N=4 mice).

Taken together, these results indicate that the SC mostly provides a nonsensory contribution to audiovisual decisions, leaving the cortex to process and integrate the sensory signals. The results of inactivations causally implicate visual cortex (VIS) in the processing of lateralized visual signals^14^, with a prominent impact on the contralateral visual sensitivity *v*_*C*_ (**Figure 7**p). Inactivations of prefrontal cortex (MOs) indicate a role in the processing of audiovisual signals, impacting both visual and auditory sensitivity from both sides, and providing strong contralateral action pressure (**Figure 7**p). Finally, consistent with our recordings in SCm (**Figure 4**b,c), the inactivations demonstrate that SC influences both Left/Right and Go/NoGo decisions, with a distinct causal role on contralateral action pressure *β*_*C*_ (**Figure 7**p).

Our findings therefore indicate that the elements of the 3-way logistic classification model have a direct mapping on distinct brain regions. In this mapping, visual and auditory^14^ areas of the cortex provide mediate the visual and auditory sensitivities (*v* and *a*); the two sides of SCm provide competing contralateral action pressures *β* that are independent of sensory stimuli; and that the prefrontal cortex integrates these signals into two decision variables *z*_*L*_ and *z*_*R*_. As for the softmax operation itself, it might be performed in some neurons of; and that finally the prefrontal cortex itself, or in a downstream integrator ^14^ in the brainstem (not in the SC), thus, performs the softmax operation, making the selection among the eligible actions (**Figure 7**q).

## Discussion

We examined the roles of the superior colliculus (SC) in a logistic decision based on auditory and visual inputs and found that the SC provides powerful additive pressure in favor of competing actions, independently of sensory inputs. The recordings revealed that the SC encodes visual and auditory stimuli mostly in separate neurons, and upcoming actions in yet other neurons. Furthermore, the inactivations revealed that rather than contributing to contralateral visual or auditory sensitivity, each side of the SC strongly promotes contralateral choices over inaction and mildly suppresses ipsilateral choices. In our task, therefore, the two sides of SC work in concert, simultaneously promoting left and right actions over inaction. These results are parsimoniously captured by a three-option logistic model in which the SC provides constant, competing action pressures that promote eligible actions relative to inaction.

This understanding of the role of the SC was made possible by the fact that the task had three possible outcomes: Left choices, Right choices, and NoGo choices. In these circumstances, the 3-option logistic classifier ^2,15,16,102^ that described the mouse behavior involved not only the usual sensory terms, but also two nonsensory additive constants *β*_*L*_ and *β*_*R*_. We termed them “action pressures” because they each promote one of the two eligible actions relative to inaction. The ability to estimate these separate action pressures proved fundamental for interpreting the results of inactivations. By contrast, a previous 2-class model ^14^, which explained the probabilities of Right vs. Left choices (ignoring NoGo choices), could only estimate a single overall bias *β = β*_*R*_ − *β*_*L*_ in favor of Right and against Left choices.

Our recordings argue against a prevalence in mouse SC of integrative computations such as audiovisual integration or sensorimotor transformation. Visual and auditory responses were mostly segregated in distinct layers, and in contrast with many reports ^21–31^ emphasizing audiovisual integration in the SC of multiple species (including the mouse^23,31^), most SC neurons were strikingly unimodal, with only 2% of all neurons being both auditory and visual. Perhaps the scarcity of audiovisual neurons in our experiments is specific to our audiovisual stimuli or to the fact that all our mice were trained in our audiovisual task (none were naïve). Additionally, some of the audiovisual responses described in earlier studies might have reflected rapid sound-evoked changes in behavioral state or perhaps even rapid orienting movements (whether executed or just planned) that were triggered by the stimuli, which can affect neural responses across the brain^120,121^.

The mostly unimodal sensory responses of SC neurons remained similar during passive presentation and during task performance. Task engagement, however, provided a general additive and multiplicative effect on the responses, which were captured by an engagement kernel. Impending actions, moreover, were often associated with increases in activity, which were captured by two action kernels. Once again, these task-related signals were largely carried by different neurons from those that encoded stimuli. This anatomical and functional separation indicated that the SC processes sensory and task-related signals largely independently.

Another clue to the role of SC came from our decoding analyses, which revealed trial-by-trial choice encoding beyond what was already predictable from the stimuli. This effect was present also in the activity from prefrontal cortical area MOs and was present only after stimulus onset. Therefore, any contribution by SCm or MOs to the choices is likely to occur during the trial, and not to be predetermined before trial onset, as had been observed in multiple brain regions^122,123^.

The unilateral inactivations further revealed that SCm provides stimulus-independent pressure for eligible actions. Unilateral SCm inactivation shifted choices away from the contralateral side and toward inaction, without altering sensory accuracy. By contrast, sensory sensitivity was greatly reduced when we inactivated visual cortex. In our task, therefore, the processing of visual signals is strongly cortical, unlike other visual tasks that may rely at least partially on SC ^20,124,125^. Analogously, sensitivity to contralateral auditory and audiovisual stimuli in our task appear to arise mostly outside the SC, because it was unaffected by unilateral SC inactivation. Therefore, the SC neurons that are visual, those that are auditory, and the few that are audiovisual do not noticeably contribute to the sensory sensitivity of the mice in the task.

The bilateral inactivations revealed that SCm provides similar pressure for all eligible actions and provided both confirmation and a reinterpretation of the classical Sprague effect. Starting with Sprague and Meikle’s early observations ^36,90^, there have been multiple reports that bilateral suppression (via inactivations or lesions) of SC resolves the effect of unilateral suppression ^39,86– 89,110,111,126,127^. To some extent, our results confirm this view, as bilateral inactivation restored balance between left and right actions and restored reaction times. However, we also found that it makes actions less likely, reducing the odds of actions over inaction. The overall reduction of action pressure was, however, slightly smaller than predicted by simply adding the reductions in contralateral action pressures, because these reductions were accompanied by weak increases in ipsilateral action pressures.

The 3-option logistic classifier explains these effects quantitatively, showing that the phenomenon arises from a rebalanced competition among three options (Left, Right, and inaction) rather than restoration of motor execution functions. Inactivating one side of SC made the contralateral action less likely relative to inaction. Inactivating the other side as well made the opposite action also less likely relative to inaction. The balance between the two actions was thus restored, but the balance between action and inaction was tilted in favor of inaction. The overall balance was set by each hemisphere strongly promoting contralateral actions and weakly suppressing ipsilateral ones. During normal function, therefore, the intact SC promotes all eligible competing actions, supporting an equilibrium across them while advantaging them over inaction.

Some of the competition between actions may take place within the SC itself. Indeed, our data suggest that while each hemisphere primarily promotes contralateral action pressure, it simultaneously slightly prevents ipsilateral actions, perhaps through mutual inhibition between the two SC hemispheres ^42,44,82,111^. This mutual inhibition would be consistent with increases in firing rate seen contralaterally when inactivating SC unilaterally^82^, suggesting local competition.

The results of cortical inactivations indicate a division of labor, where SC provides action pressure, visual cortex provides visual signals, and prefrontal cortex integrates audiovisual signals and provides some action pressure. Indeed, the unilateral cortical inactivations gave markedly different results from the unilateral inactivations of SC, as they severely degraded contralateral sensory sensitivity. These inactivations revealed that in this task the causal role of visual cortex (VIS) is visual, and the causal role of prefrontal cortex (MOs) is audiovisual. In other respects, however, inactivations of MOs more closely resembled inactivations of SC: unilateral MOs inactivations slightly reduced contralateral action pressure, a reduction that might not be as strong as the reduction seen with SC inactivation but was nonetheless present. Moreover, when the inactivations of SC were bilateral, they did slightly affect sensory sensitivity.

These findings rule out the possibility that the SC is a station in a single pathway proceeding downstream from the prefrontal cortex. Indeed, were SC part of an obligatory path, inactivations there would have affected not only contralateral action pressure but also stimulus sensitivity. Moreover, bilateral inactivations would have abolished movements entirely or at least slowed their execution. We can thus rule out the view of SC as a simple motor trigger ^37,47,52^ and reinforce the view that the SC contributes to decision processes via action or target selection ^54,65,70^.

Our results, however, leave open two possibilities for the destination of SC signals. One possibility is that action pressures provided by SC are sent up to the prefrontal cortex (perhaps via thalamic areas), which integrates them with sensory evidence from sensory cortices, and perhaps resolves the competition between eligible actions. This view would be supported by ample evidence for upstream projections of SC ^54,55,80^ and by observations that SC inactivation modulates prefrontal cortex ^82^. In this circuit, the superior colliculus proposes available actions, while the cortex resolves the competition based on sensory evidence. Alternatively, or additionally, SC may project to a downstream integrator that also receives input from prefrontal cortex. Finally, the basal ganglia may also play a major role^44,46–48,50,80^, which we have not explored in our study.

Taken together, these results demonstrate that the superior colliculus plays a major role in defining the space of eligible actions and in promoting the competition between those actions. By promoting all competing eligible actions, the colliculus may make it much more likely for one of the eligible actions to win the competition over ineligible ones, which include inaction. More generally, the discovery that the colliculus provides action pressure independently of external stimuli agrees with the view that decision-making circuits are fundamentally organized around competing motor affordances^128,129^. Our results define the computations underlying this competition and the major role that the colliculus plays in supporting them.

## Supplementary material

**Supplementary Figure 1.**
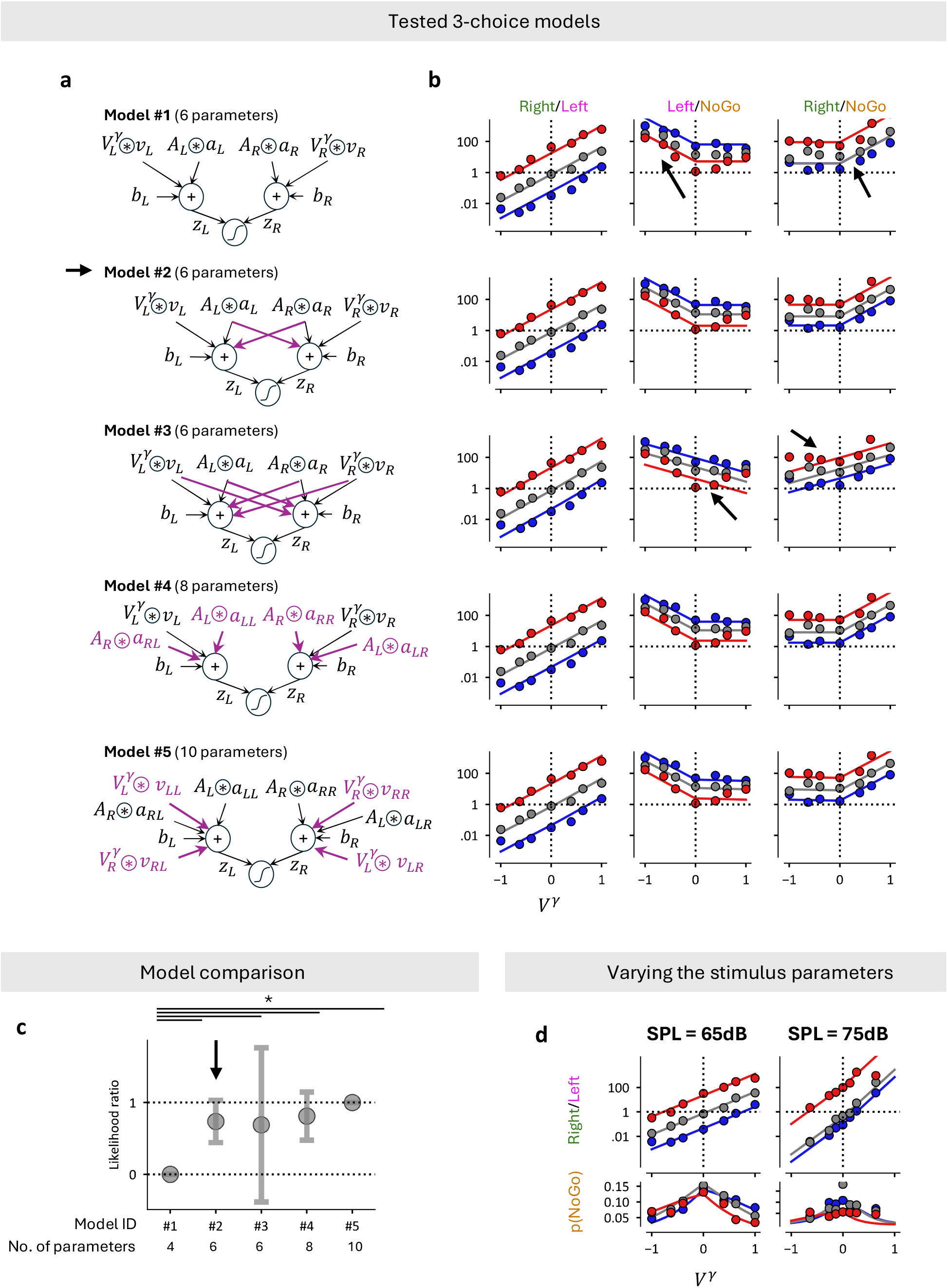
Model selection. **(a)** Schematics of the 5 different models we tested. Model 1 is a lateralized sensory model where the decision variable z_L_ is only influenced by left stimuli and vice versa. Model 2-3 are symmetric models, where each decision variable is influenced by both left and right stimuli but with the same weight. In contrast, in models 4-5 each stimulus can be weight differently for z_L_ vs z_R_. **(b)** Fits of each model, represented as the odds of each choice combination. **(c)** Model performance on the test set normalized between the simplest model (Model 1) and the model with the most parameters (Model 5). Line is the average, and the shaded area indicates 95% confidence interval across subjects. Model performance significantly differed from all other models, but otherwise, we could not distinguish a systematic difference between models 2-5. (repeated measures ANOVA across 5 models within subjects (p<0.001), followed by a post-hoc comparison using paired t-test with Bonferroni correction.) **(d)** Visual comparisons of model 3 performance across two different cohorts of mice when different stimulus intensities were used. Left: mice where sounds were played at 65dB and we used 10,20 and 40% contrast levels for visual stimuli and left: sounds were played at 75 dB and we used 5,10 and 25% contrast levels as visual stimuli.

**Supplementary Figure 2.**
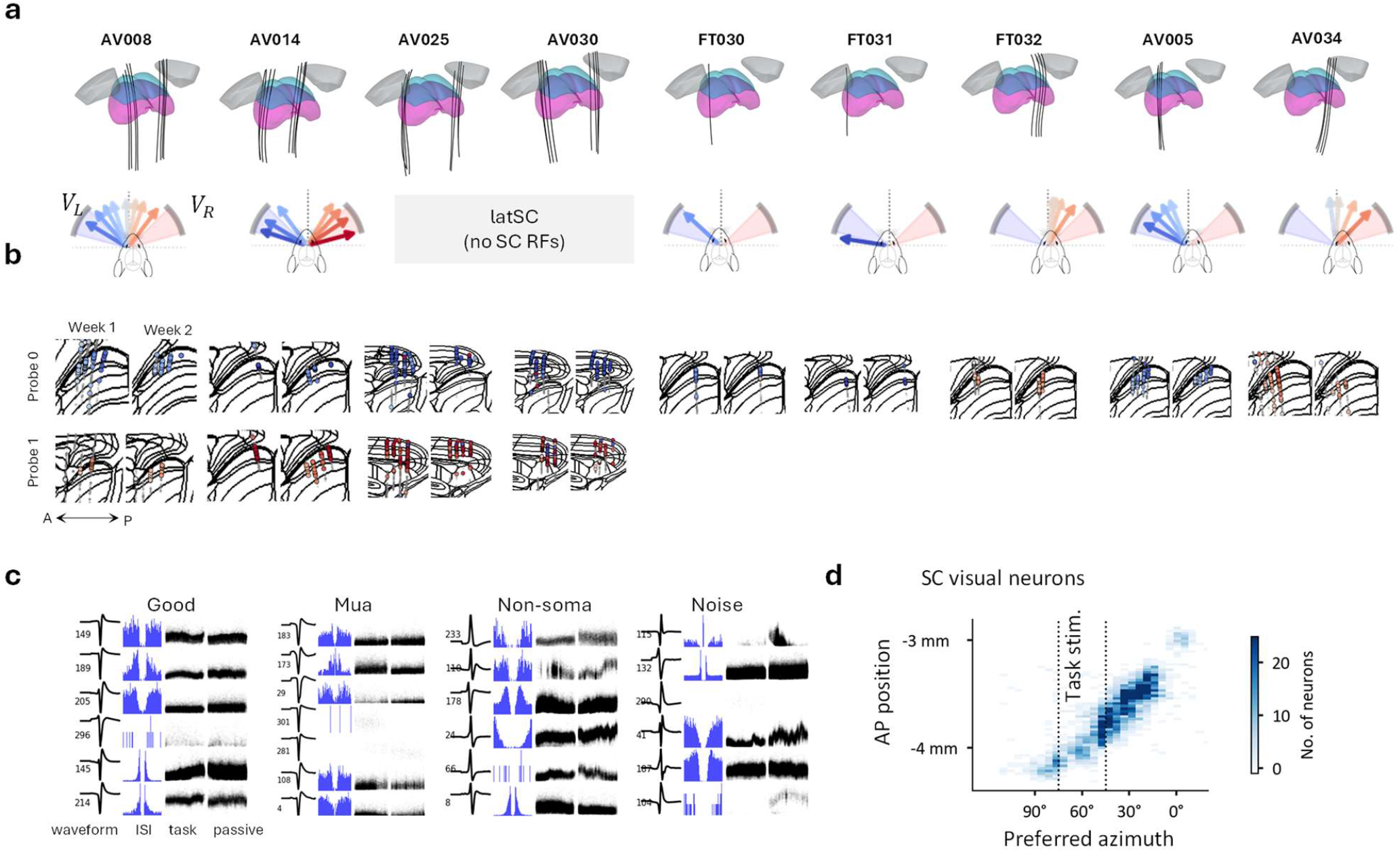
Electrophysiology quality control. **(a)** (top) Location of shank insertions into SC registered to the Allen atlas (25 µm) per each mouse. (bottom) Preferred visual azimuth on each shank in each mouse, calculated by fitting receptive fields from sparse noise stimuli. **(b)** SC Receptive fields per neuron on 2 consecutive weeks after implantation. Each pair corresponds to the same mouse as in (b). Some mice had two probes inserted (probe0 and probe1). **(c)** Example units that were classified (from left to right) as good neurons, multiunit, non-somatic and noise. Units were clustered using pyKilosort ^130^ and curated by Bombcell ^131^. Each row is a unit, and for each unit we show (from left to right) the waveform, the interspike-interval (ISI) histogram and a subsampled trace of the amplitude of spikes across time during the entire task and passive session (~1 hour each). **(d)** Density of preferred azimuths (multiunit activity) vs AP location (relative to Bregma). Dashed line: task stimulus location, showing task stimuli are represented ~4mm posterior to Bregma.

**Supplementary Figure 3.**
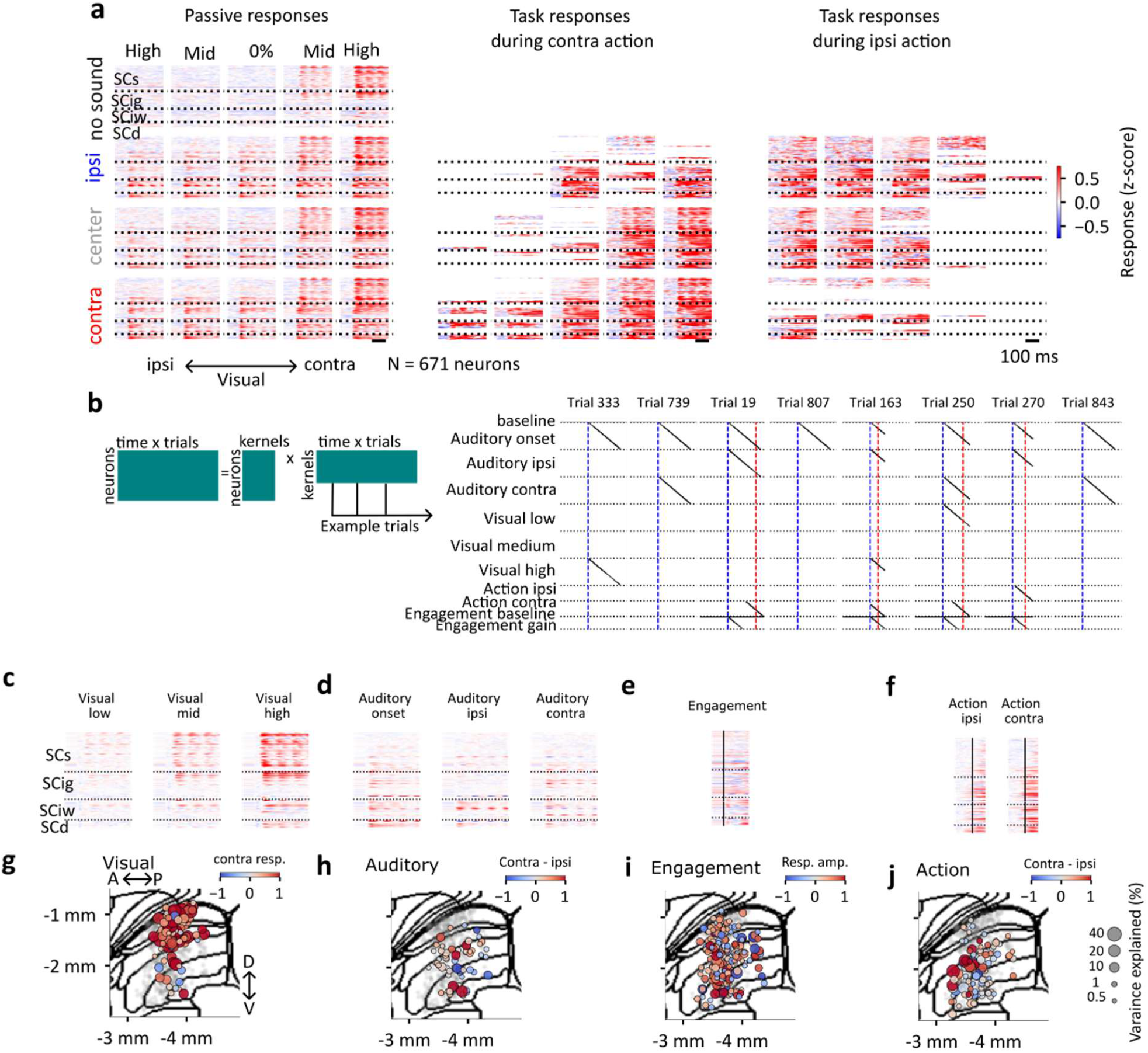
Further details of fitting the encoding model on SC data. **(a)** Responses of SC neurons (n = 671), sorted by depth during passive stimulation and contra- and ipsilateral actions (left to right). Matrices show responses across visual contrast (x-axis) and auditory stimulus location (y-axis), as in **Figure 2**f-i. Missing lines indicate that animals did not make a particular choice frequently on that session preventing us to show trial averages. During task, the ‘no sound’ condition was not presented. **(b)** Cartoon of the encoding model. We predicted neural activity over trials (−0.2 s to 0.55ms or until 0.1s after reaction time) using Ridge Regression using temporal basis functions, i.e. kernels. We predicted neural activity simultaneously during passive replay and task execution. Example trials and the supported kernels in the given trial are shown in the design matrix (trials selected randomly). Blue vertical time indicated stimulus onset, red line indicates action initiation. Black lines indicate ones where the kernel is supported, else the matrix is filled with zeros. **(c-f)** Visualization of visual (c), auditory (d), engagement (e) and action (f) kernels learnt for a given neuron (rows), sorted by depth in the SC. As neurons were z-scored before fitting, kernel size approximates the response magnitude. Task kernel is the sum of engagement baseline and engagement gain kernels. All kernels are triggered on stimulus onset except the action kernels, which are triggered on action initiation. **(g)** Visual neuron locations (AP-DV plane, coloring dots with >0.5% variance explained by visual kernels). Dot size indicates % variance explained. Color indicates response magnitude (red – excited, blue – suppressed) **(h)** Auditory neuron locations colored by tuning direction (*blue*: ipsi, *red*: contra) **(i)** Engagement modulated neurons colored by modulation amplitude **(j)** Action modulated neurons colored by their preferred action direction (*blue*: ipsi, *red*: contra).

**Supplementary Figure 4.**
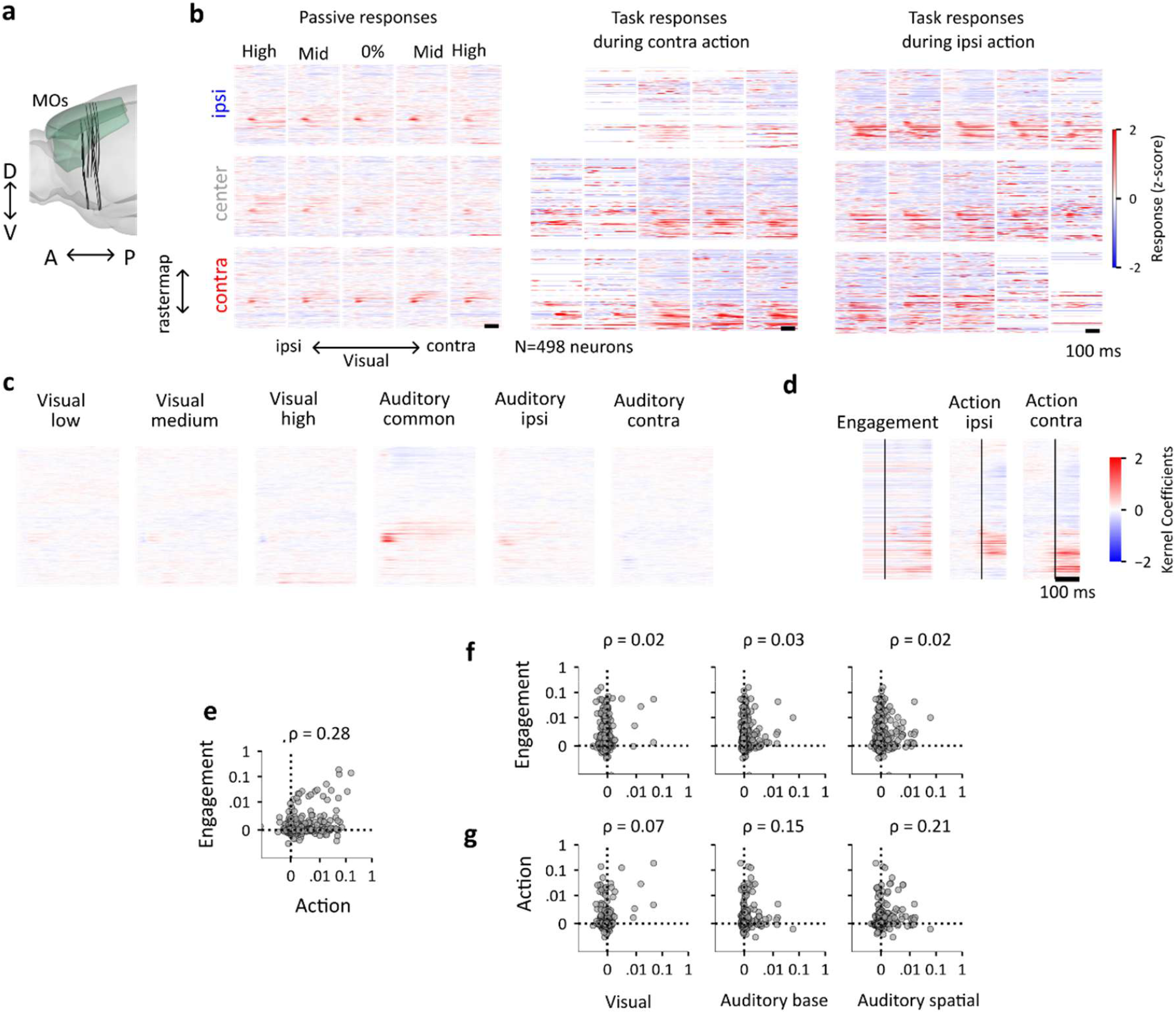
MOs activity characterized by the encoding model. **(a)** Anatomical position of MOs recordings (16 shanks, 4 probes, 3 mice). **(b)** Responses of MOs neurons (n=498) sorted by Rastermap during passive stimulation and contra and ipsilateral actions, like Supplementary Figure 3a. **(c)** Sensory kernels for neurons in (b). We applied the same model to the MOs data as we presented for SC in Figure 2-3. **(d)** Engagement and action kernels for neurons in (b). **(e)** Pairwise comparison of variance explained by engagement versus action modulation in MOs neurons. **(f)** Pairwise comparison of variance explained by sensory versus engagement kernels in MOs neurons. **(g)** Same as (e), but for sensory versus action kernels.

**Supplementary Figure 5.**
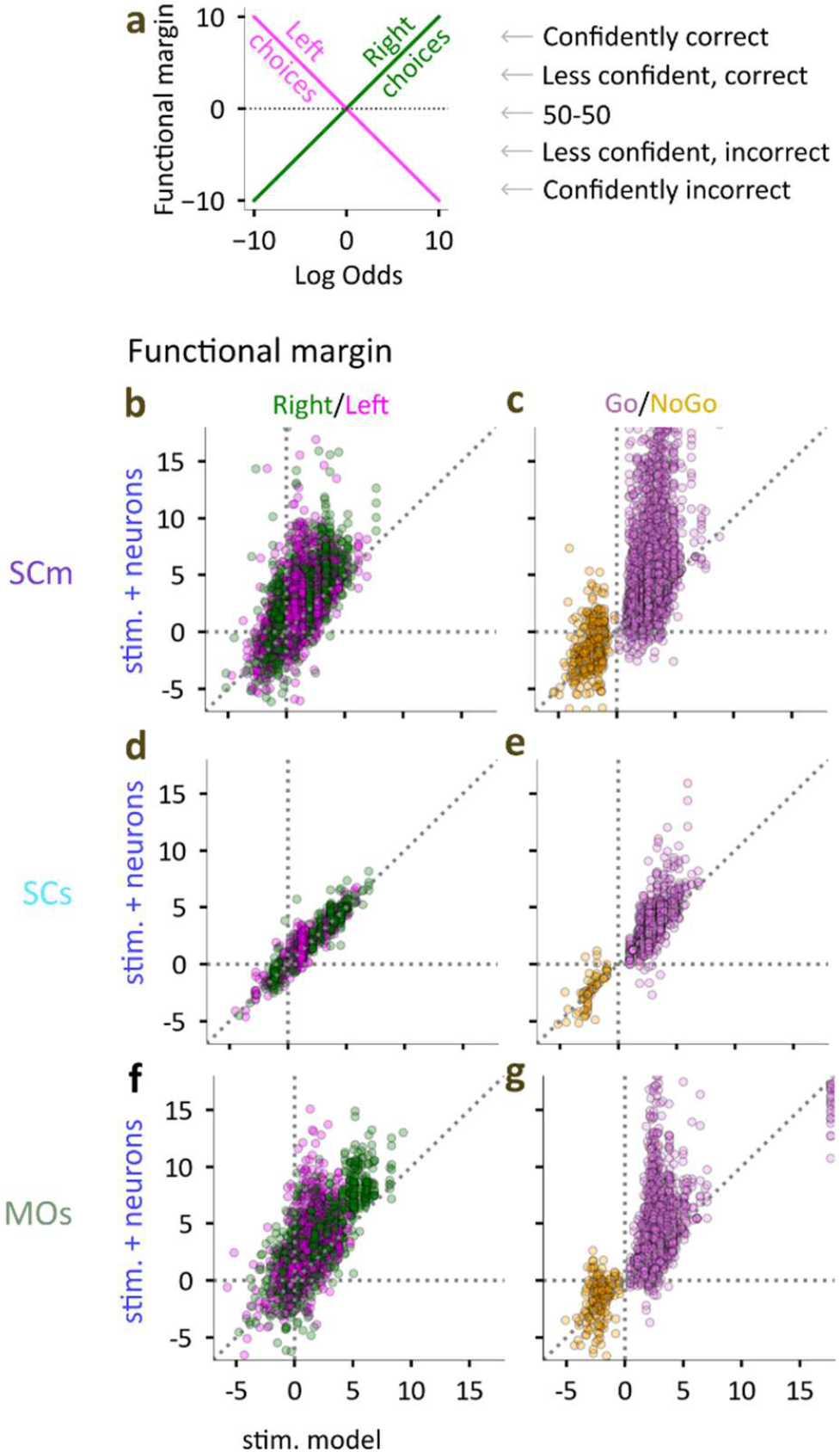
Further diagnostics of the neural decoder. **(a)** To assess model performance, we multiplied the predicted log-odds by the actual outcome (−1 or +1) to obtain the functional margin. Positive values denote correct predictions, negative values denote errors, and magnitude represents model confidence. We made this calculation for both Left/Right outcomes (as shown here) and Go/NoGo outcomes. **(b)** Left vs Right decoding using SCm activity, plotting the functional margin for the stimulus-only vs. the extended (stimulus + neural) model. Each dot indicates a trial, colored according to behavioral outcome. (4,128 trials, 26 sessions in 9 mice) **(c)** Same, for Go vs NoGo decoding (4,547 trials, 26 sessions in 9 mice). **(d-e)** Same as b-c, for SCs neurons. (n = 1,046 trials, 6 sessions, 4 mice) **(f-g)** Same, for MOs neurons (2,051 Right vs Left trials, 2,254 Go vs NoGo trials, 19 sessions in 3 mice).

**Supplementary Figure 6.**
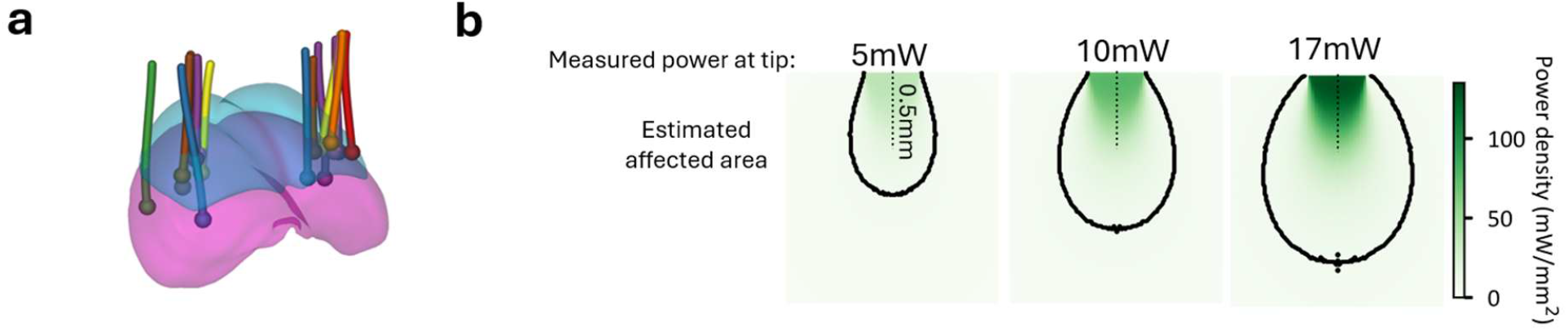
Additional details of SC inactivation experiments. **(a)** Cannulae locations in SC. **(b)** Monte-Carlo simulations of light spread (565nm wavelength) in tissue at various tip powers used for inactivation^132^, showing the power density around a 400µm tip. The contour of the area where the power crosses the EPD_50_ of eNpHR3.0 is shown as a black line.

**Supplementary Figure 7.**
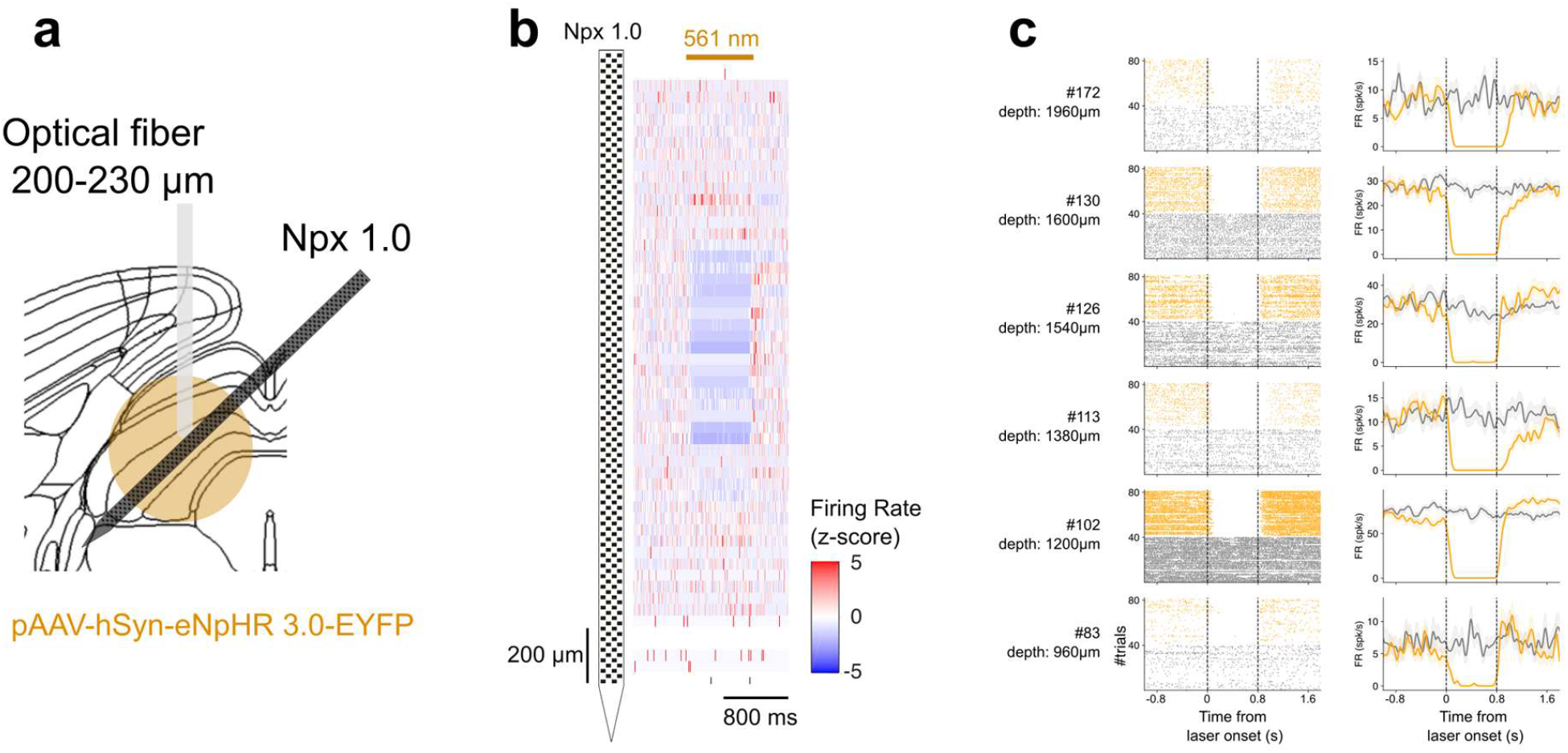
Validation of optogenetic SC inactivation near the cannula. **(a)** Schematic of the experimental approach. A viral vector expressing halorhodopsin (pAAV-hSyn-eNpHR 3.0-EYFP) was injected into the left superior colliculus (SC) to enable optogenetic inhibition. An optical fiber (DORIC, C60, 200–230 µm diameter) was implanted above the intermediate and deep layers of the left SC for light delivery. Neuropixels 1.0 probe was used for electrophysiology recordings performed 4 weeks after viral injection and inserted from the right hemisphere. **(b)** Heatmap of z-scored average firing rates (bin size 50 µm, 40 trials) plotted as a function of depth during passive laser stimulation (Omicron 561nm laser, 800 ms pulse, 10mW measured at the optical fiber tip). Units recorded near the optical fiber tip show significant suppression of activity (*blue*), confirming local inactivation of SC neurons near the cannula. **(c)** Single-unit examples recorded near the cannula at varying depths (same session as in (b). Raster plots (left) and peri-stimulus time histograms (right) are shown for baseline (no stimulation, *gray*) and passive laser stimulation (*orange*) conditions. The firing rate was reliably and substantially reduced during laser trials. Dashed lines indicate laser onsets and offsets.

**Supplementary Figure 8.**
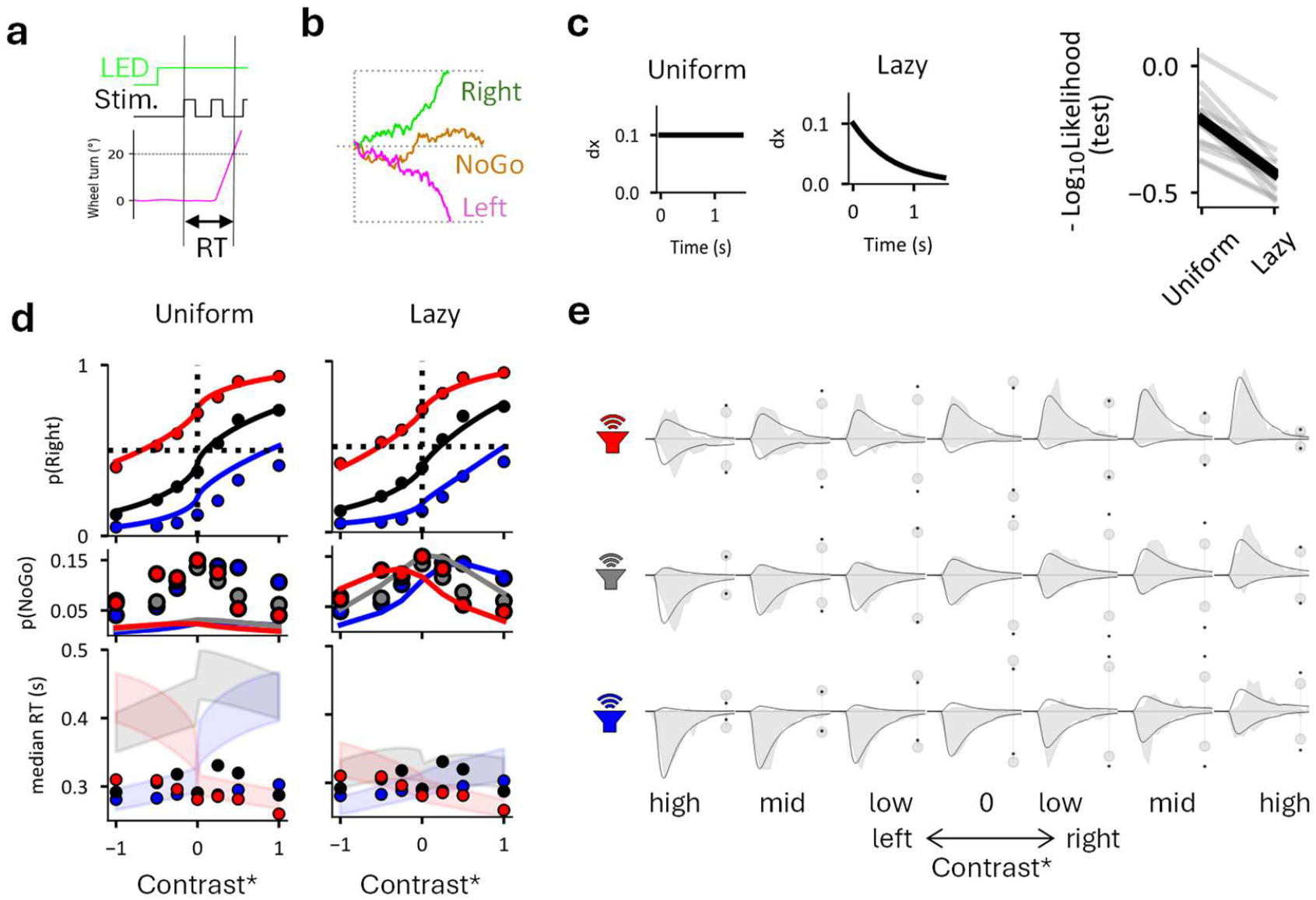
Modeling the audiovisual task with a modified DDM. **(a)** Temporal trial structure during inactivation trials. Reaction times are defined as the interval between decision completion and first stimulus onset. **(b)** Three example simulations of the drift–diffusion process showing left, right, and NoGo outcomes. Evidence (x, y-axis) accumulates over time (x-axis) via incremental updates (dx) determined by the drift rate and Gaussian noise. The decision terminates when evidence reaches the upper (right) or lower (left) bound. If neither bound is reached within the response window, the trial is classified as a NoGo. **(c)** (left) Drift-rate dynamics as a function of time since stimulus onset for the two models compared: uniform drift and lazy drift, in which drift rate is scaled by an inverse gain function. (right) Model comparison between uniform and lazy drift models quantified by negative log_10_-likelihood (lower values indicate better fit; n = 14 mice). **(d)** Comparison of diagnostic plots for the uniform and lazy drift models (left vs. right columns). From top to bottom: fraction of right choices as a function of visual contrast (x-axis) and auditory stimulus location (color), fraction of NoGo choices, and median reaction times. Dots indicate data and lines indicate model predictions. In the reaction-time plot, shaded areas represent the 45th– 55th percentiles of the predicted distribution (n = 10,598 trials, 754 trials each from 14 mice). **(e)** Data (*gray*) and predictions (*black*) from the lazy drift model across stimulus conditions (visual contrast × auditory location). Each panel shows reaction-time distributions for right (upper) and left (lower) choices and the relative fraction of NoGo choices (plotted on both sides as dots). Data are identical to those in panel (d).

**Supplementary Figure 9.**
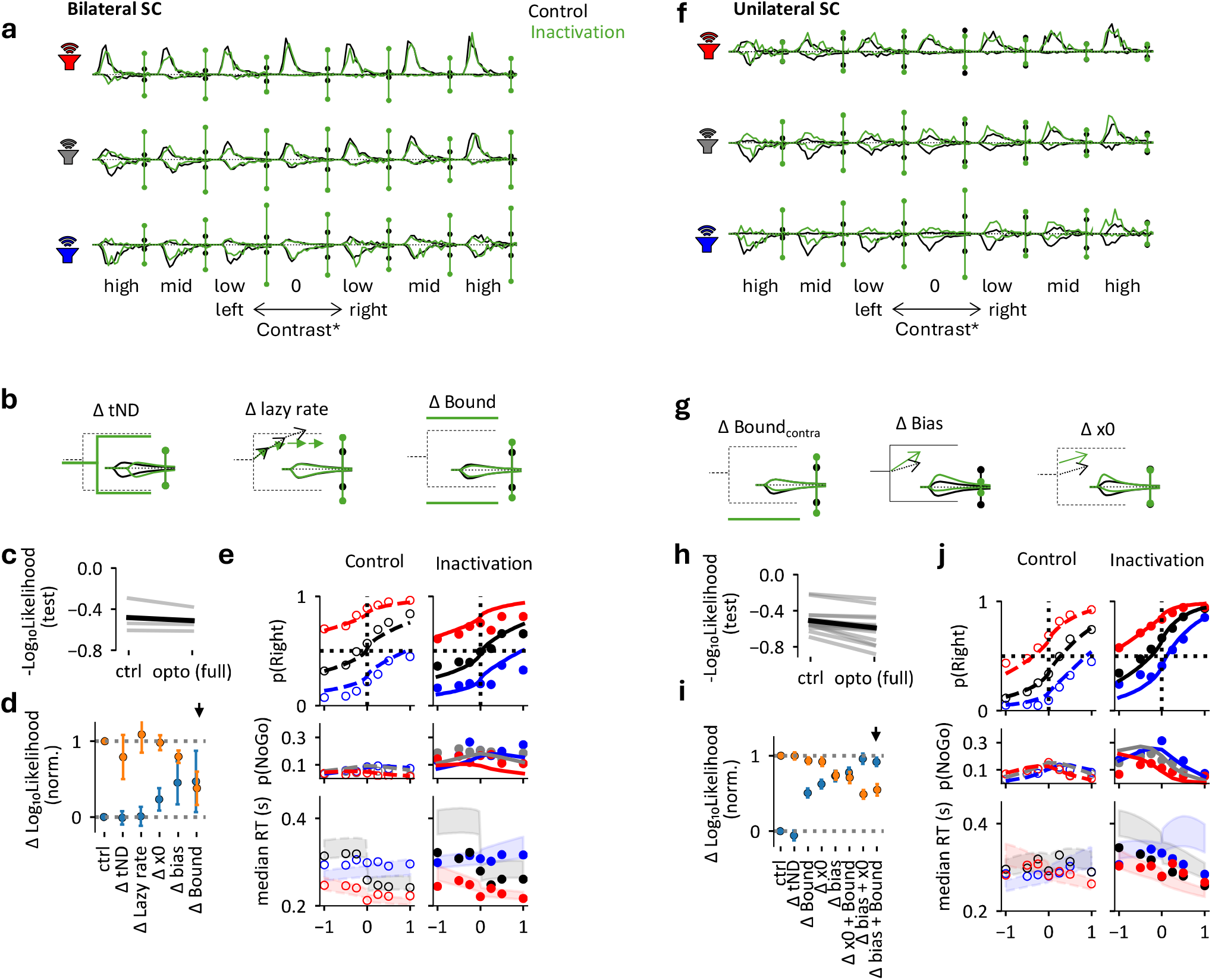
Modeling SC inactivation effects with the DDM. **(a)** Reaction-time (RT) distributions and NoGo fractions shown as in **Supplementary Figure 8**e for control (black, 12,970 control trials, 4 mice) and bilateral SC inactivation trials (green; n = 3,558 trials). **(b)** Illustrations of candidate parameter changes explaining altered NoGo rates after bilateral inactivation: changes in non-decision time (ΔtND), lazy rate, and an expanding bound. Insets show simulations of blank trials (*black*: control; *green*: inactivation). An increase in tND shifts reaction times uniformly along the x-axis with minimal effect on NoGo rates. An increase in the lazy rate primarily increases the fraction of NoGo trials, whereas an increased bound slightly slows reaction times and increases NoGo choices. **(c)** Log_10_Likelihood of the control model and the extended model jointly fitted on control and inactivation trials. The extended model allowed the nondecision time (ΔtND), the lazy rate, the starting point (Δx0) the constant drift bias (Δbias) and the bound to change by inactivation. **(d)** Likelihood ratio test assessing each parameter’s contribution to model fit. Blue dots indicate log-likelihood gains relative to the control model (mean ±s.e.); orange dots indicate losses relative to the extended model. Parameters are ordered by increasing likelihood gain. **(e)** Diagnostics (as in **Supplementary Figure 8**d) of the ΔBound model on control (left column) and inactivation trials (right column). **(f-j)** Same analyses as panels (a-e) for unilateral right-hemisphere SC inactivation (n = 2,813 opto trials, 9,217 control trials, 17 hemispheres, 10 mice). Simulations illustrate parameter changes producing lateralized effects, including asymmetric bound changes (contralateral, ΔBound), drift-rate bias (Δbias), and starting-point shifts (Δx0) and their combinations. A contralateral increase in bound (ΔBound) slightly increases reaction times while producing a large increase in NoGo choices. Changes in drift-rate bias (Δbias) strongly alters choice ratios with minimal effects on median reaction times. Changes in the starting point (Δx_0_) produce strong asymmetric effects on reaction times while also altering choice fractions. In panel (j) we plotted the overall best fitting model for unilateral inactivation, which simultaneously allowed for a bias and asymmetric Bound change.

**Supplementary Figure 10.**
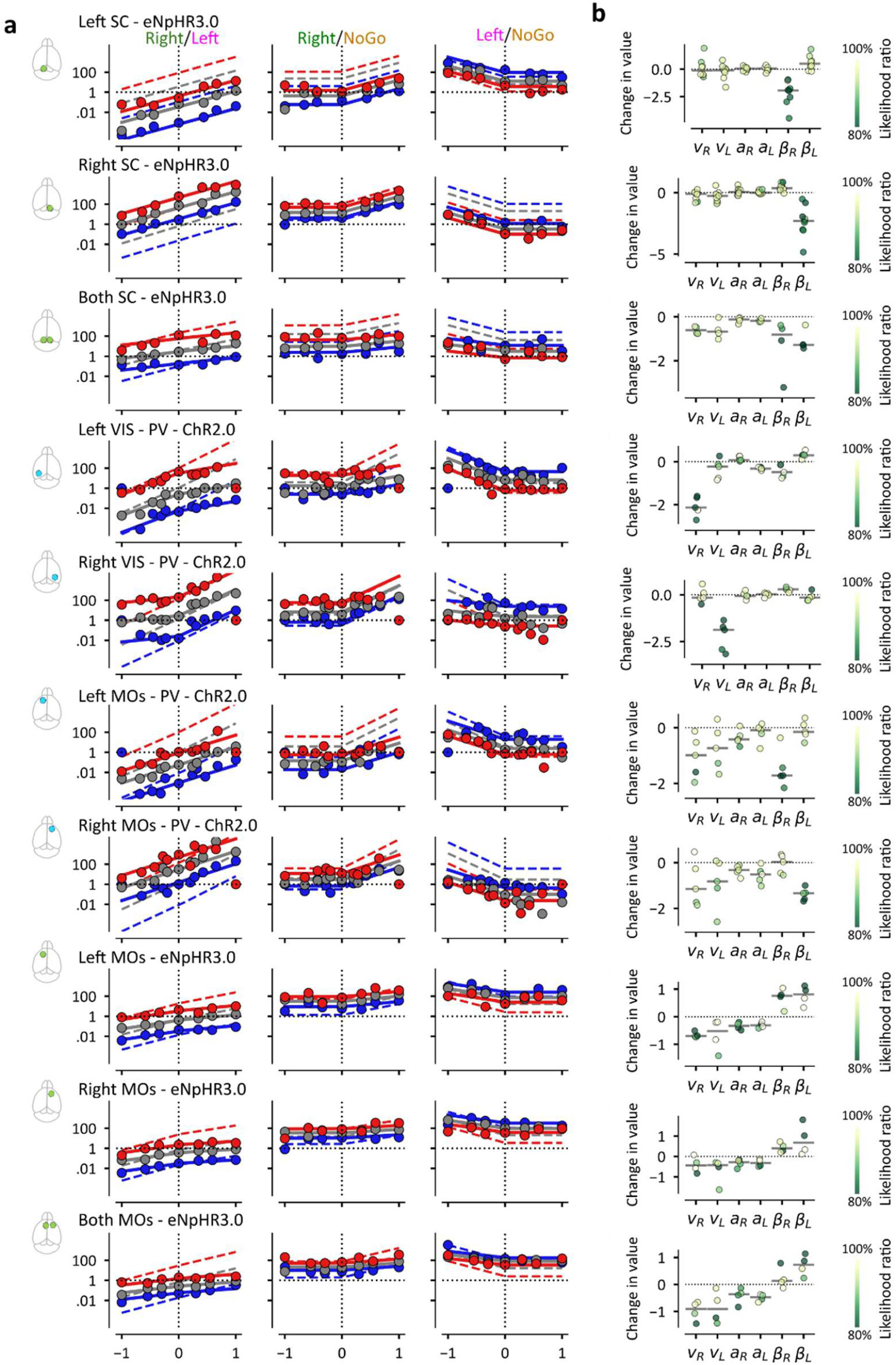
Further details of psychometric fits across brain regions. **(a)** Changes in model fits by various inactivation conditions: from top to bottom: left SC, right SC, both SC using eNpHR3.0, left VIS and right VIS using PV-ChR2.0, Left MOs and Right MOs using PV-ChR2.0. PV-ChR2.0 data from Ref. ^14^. Left MOs, Right MOs, Both MOs inactivated with eNpHR3.0. **(b)** Changes in parameter values by each inactivation. Dots indicate mice and are colored by the likelihood ratio value relative to the extended model.

## Acknowledgements

We thank Carolina Quadrado for assistance with histology and Bex Terry for assistance with animal husbandry and the monitoring our animals’ health, and Charu Bai Reddy for laboratory management and surgical training. The authors also acknowledge the use of the UCL Myriad High-Performance Computing Facility (Myriad@UCL), and its associated support services. During the preparation of this work, we used Gemini or Claude frequently to assist in coding and occasionally to suggest edits to the text. We reviewed and edited the output as needed and take full responsibility for the content of the published article.

This work was supported by the Wellcome Trust (Investigator Award 223144/Z/21/Z to MC and KDH) and by BBSRC (award BB/T016639/1 to MC). F.T. was supported by the Sainsbury Wellcome Centre PhD program. M.C. holds the GlaxoSmithKline/Fight for Sight Chair in Visual Neuroscience.

## Code and data availability

Code used to preprocess data is available at github.com/cortex-lab/PinkRigs. Code to 3-choice fit behavioral and neural models is available at github.com/takacsflora/NeuroLogit. Code to reproduce figures and neural data will be shared upon publication.

## Author contributions

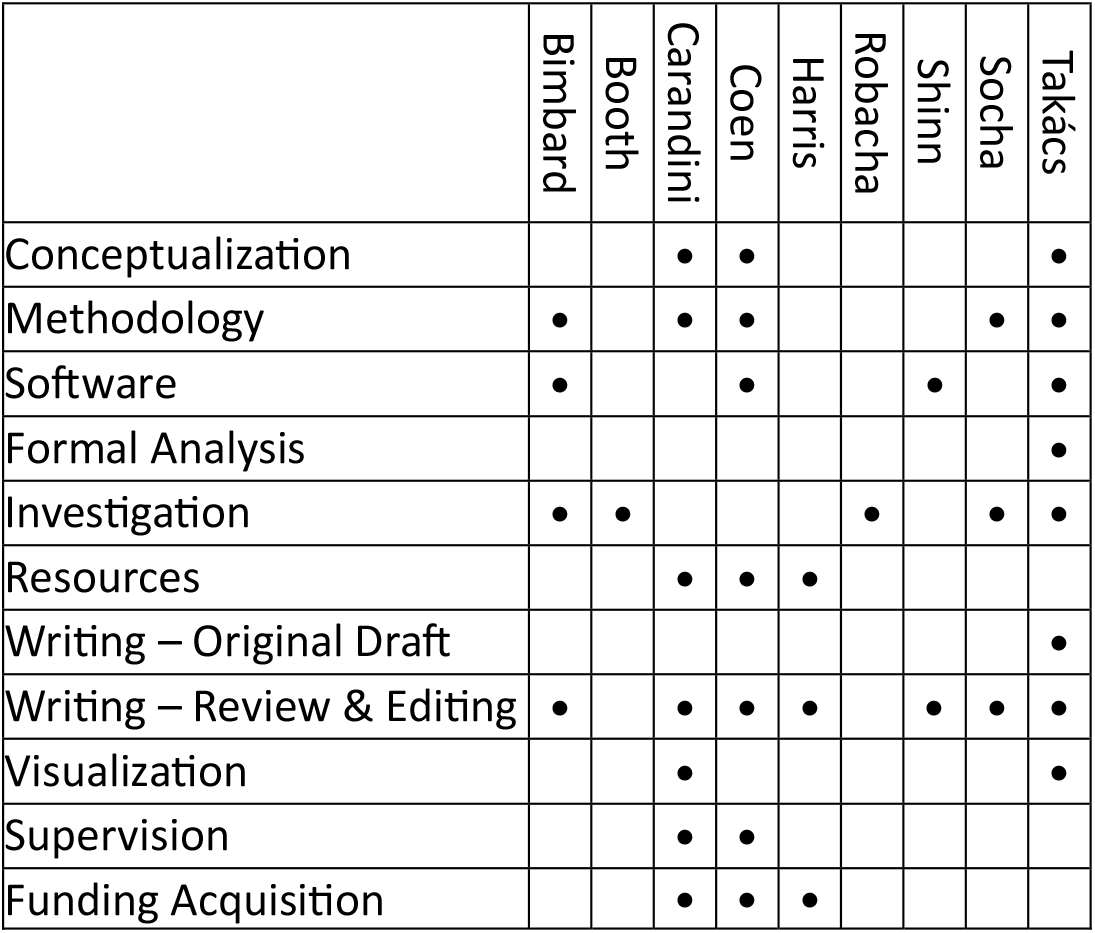

## Methods

Experiments were conducted according to the UK Animals Scientific Procedures Act (1986) and under personal and project licenses released by the Home Office following appropriate ethics review.

### Mice

Mice (18-45 weeks) were Cdh23-positive (hearing loss corrected) with the C57BL/6N background ^133^. For SC recordings, we used 7 male and 2 female mice (4 implanted bilaterally and 5 unilaterally). For MOs recordings, we used 3 female mice (2 with one inserted probe and 1 with two probes). For optogenetic inactivation of the SC, we used 2 female and 8 male mice (7 implanted bilaterally and 3 unilaterally). Only 4 of the 7 mice implanted bilaterally were also inactivated bilaterally, because the two cannulas were too close to each other for two patch cords to fit. For MOs inactivations with eNpHR3.0, we used 3 female and 1 male mice, all implanted (and when needed inactivated) bilaterally. Mice were assigned to our study independently of sex once confirmed Cdh23-positive and above the minimum implant weight threshold.

### Surgeries

All mice underwent two surgeries under isoflurane anesthesia. First, we implanted a steel headplate as described previously^14^. In the second surgery, we implanted either optical cannulae or Neuropixels probes.

#### Optogenetics preparation

To inactivate SC or MOs, we used pAAV-hSyn-eNpHR3.0-eYFP (Addgene plasmid #26972, gift from Karl Deisseroth). We made one or two 1 mm craniotomies using a biopsy punch (for uni- or bilateral prep) and injected 110-140 nl virus diluted in ACSF at 3 different depths separated by 150µm. We used a 2.5*10^12^ GC/ml titer for SC injections 5*10^12^ GC/ml titer for MOs injections. After injection, we placed a cannula at the top of the injection spots and cemented it to the skull (Doric Lenses: MFC_400/430_4mm_MF1.25_C60, for SC, MFC_400/430_mm_MF1.25_C60 for MOs).

#### Neuropixels implants

To record in the SC, we used Neuropixels 1.0 or 2.0 probes implanted chronically ^104,105^. We implanted probes with a non-recoverable or recoverable method as described previously, after mice were already trained in the task. Briefly, for the non-recoverable method, the Neuropixels probe was inserted and cemented onto the skull during the second surgery through a small craniotomy^134^. For the recoverable method, we implanted the probes with a two-module construct, where we cemented the docking module to the skull, not the probe itself ^103^. This allowed us to recover the probes as we ended the experiments (typically due to low neural yield).

##### Histology

We perfused mice transcardially and post-fixed brains in 4% formalin-PBS for 24 h. We imaged brains with two-photon serial tomography ^135^ and aligned them to the 25 µm Allen Mouse Brain Atlas ^136^ using *brainreg*^*137*^. We traced cannula and probe tracks manually in atlas space using *brainreg-segment*^*138*^ and visualized tracks using *brainrender* ^139^.

### The audiovisual decision-making task

We presented all stimuli using RIGBOX ^140^.

#### Visual stimulation

Visual stimuli were displayed on three screens (Adafruit, LP097QX1) positioned ~11 cm from the mouse, covering ±135° azimuth and ±37.5° elevation. We equalized luminance viewing angle with Fresnel lenses (Wuxi Bohai Optics, BHPA220-2-5) and a diffusing film. The screens refreshed at 60 Hz.

##### Receptive field mapping

To map receptive fields, we displayed white squares of 7.5°-size on a black background. The location of white squares was selected randomly from a spatially uniform distribution. We updated the selection at 20 Hz and presented each white square for 1/6 s.

##### Task stimuli

In the task, we presented checkerboards at 60° azimuth. Each pattern contained 4 × 10 squares (7.5° each), so the visual stimulus size was 30° wide 75° tall. We paired contrasts of 5, 10, 25% or 10, 20, 40% with 65 dB or 75 dB auditory stimuli and grouped them into low (0.25), mid (0.5), or high (1.0) contrast for analysis.

#### Auditory stimulation

Seven loudspeakers (102-1299-ND, Digikey) were equidistantly placed between ±90° azimuth just below the screen. The speakers were driven with an internal sound card (STRIX SOAR, ASUS) and a custom amplifier as described before^14^. To compensate for the unique frequency response of each speaker, we recorded 2-20 kHz white noise burst using a calibrated microphone placed at the mouse’s approximate location (GRAS 40BF 1/4” Ext. Polarized Free-field Microphone) and constructed a compensatory filter that ensured equal loudness across the power spectrum.

For the task, we used 8–16 kHz pink-noise bursts. We applied an additional random filter on each trial to prevent mice from identifying the speaker from residual frequency structure; otherwise, small spectral differences could serve as location cues. Sound levels were 65- or 75-dB SPL, and we pooled these conditions.

#### Electrophysiological recordings

All Neuropixels recordings were obtained with SpikeGLX^141^. During recordings, probes were grounded to the rig and referenced through the tip. For 2.0 probes, we used vertical channel configurations for visual receptive field mapping and horizontal configurations during task recordings to maximize SC yield. We always obtained recordings during task performance first, then during passive replay. Occasionally, we also obtained receptive field maps to keep track of probe location.

#### Optogenetic inactivation

Cannulas were coupled to an LED (M565F3 Fiber Coupled LED, Thorlabs) using a patch-cord (FP400URT, NA=0.48, Thorlabs), ensuring that no light escapes into the visual field of the mouse. Power was controlled by an analog pulse (LEDD1B Driver, Thorlabs). We calibrated light output with a flat-tip cannula that matched the implanted optics.

We used powers of 5,10,17 mW total output (mapping to 40, 80, and 135 mW/mm^2^ power density). Some mice did not receive the lower powers, and one mouse received 15mW as the highest inactivation power instead of 17mW. We estimate the decay of power intensity in tissue using Monte Carlo simulations ^132^, assuming similar propagation from flat and conical tips in aqueous tissue. To estimate the affected area (**Supplementary Figure 6**b), we used the 5 mW EPD_50_ threshold for eNpHR3.0^142^.

#### Task structure

We adapted the audiovisual localization task from our previous study ^14^. During training, mice received a reward by turning the wheel so that the stimulus moved from its location to the center, an action that mirrors an orienting movement ^15^. After training, the same movement was rewarded, but the stimuli remained in place until the action was concluded. A rotary encoder measured wheel movement (Kubler 05.2400.1122.0360). Mice initiated trials by holding the wheel for 100–250 ms (exponential distribution). Stimulus pulses lasting 50 ms were delivered at 8 Hz until the mouse made a choice. Stimuli appearing on the left were associated with clockwise wheel turns (Left choices), and right stimuli were associated with anticlockwise wheel turns (Right choices). In visual, auditory and coherent trials, correct choices were always rewarded. In conflict trials (10% of the trials) and blank trials, choices were rewarded randomly.

We introduced three main changes relative to the original task^14^: (1) we used checkerboard visual stimuli; (2) we omitted the open-loop period, and the go cue; and (3) we fixed stimulus positions after training, so stimulus movement no longer correlated with wheel movement.

We trained mice using a standardized protocol ^11^ Difficulty increased when mice met performance criteria (≥200 trials/session, ≤60% bias, ≥70% correct). We gradually reduced reward size, decreased the response window, and increased wheel gain to obtain more trials. We implanted optical fibers or probes only after full training.

During recordings and inactivation, we presented auditory, visual, coherent, and conflict trials in a 3:3:2:1 ratio. Fully trained mice had 1.5 s to respond. The minimum inter-trial interval (ITI) was 2 s. Incorrect trials triggered a timeout (up to 3 s). We repeated trials if mice made incorrect choices on the easy trials, and we excluded these trials when computing performance.

##### Inactivation trials

We inactivated SC or MOs on 33–37% of trials. We did not repeat inactivation trials even if mice made an incorrect choice. We turned the LED on after the quiescent period and before stimulus onset and randomized the LED–stimulus interval (100-200 ms) to avoid creating timing cues. Mice usually continued holding the wheel until stimulus onset on inactivation trials. The LED stayed on for 1.5 s and ramped down over 400 ms (total 1.9 s).

We also added LED-only trials (4% of all trials) in a subset of sessions. These trials enforced quiescence but contained no auditory or visual stimulus. We turned on the LED in 50% of these trials to measure movement effects driven solely by optogenetic activation.

We inactivated either the left, right or bilateral hemisphere on a given day. In most SC mice we sequentially varied the location of inactivation on each day (let-right-bi), in MOs inactivations and with the remaining SC-implanted mouse, we varied the location randomly by throwing a die before every session.

#### Measuring timings in the task

We recorded the timing of stimulus presentation, optogenetic inactivation, and wheel movements using a common DAQ (National Instruments, BNC-2090A). We also synchronized the electrophysiological recordings to this common DAQ, sending a randomized binary signal to both the DAQ and the IMEC acquisition card.

The average temporal difference in the presentation of visual and auditory stimuli was 13±7 ms, where visual display changes typically delayed relative to auditory stimulus onset. To calculate reaction times, we took the time of the stimulus that was first turned on during each trial.

To identify reaction times used for the drift-diffusion model, we used the time when the mouse brought the stimulus to the middle of the screen, i.e. the angular shift of the wheel turn reached 20°. To identify action onset times for the neural data analysis, we used first timepoint where average velocity exceeded 0.2° over a 50 ms forward-sliding window after periods of inactivity, as described previously ^15^.

### Behavioral data analysis

Prior to fitting behavioral models, we excluded trials were repeated after an incorrect choice. We also excluded NoGo trials that occurred in sequences of more than three, to ensure the model was fit only to periods where the mouse was actively engaged in the task.

For graphical purposes, whenever we plotted the log_10_-Odds of choice pairs we added one trial per each possible stimulus and choice combination to avoid infinite odds.

#### The 3-option model

We used multiclass logistic regression to predict the probability of each outcome across the three choice options: {*R, L, NoGo*} (Refs. ^15,16^). In this framework, the probability ratio of Left vs NoGo and Right vs NoGo choice pairs are determined by decision variables *z*_*L*_ and *z*_*R*_ according to Equation 1, i.e.

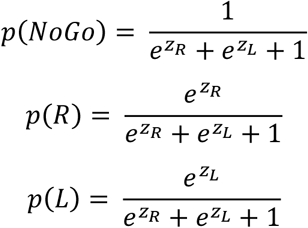

In most figures, instead of plotting the probabilities directly, we plot the log odds of the probability ratios ^2^:

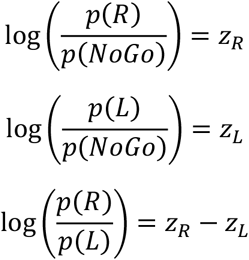

And the odds of Go vs NoGo can also be calculated as:

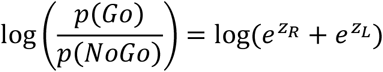

To predict *z*_*L*_ and *z*_*R*_ we used stimulus-dependent parameters (*v*_*R*_, *v*_*L*_, *a*_*R*_, *a*_*L*_) and the intercepts (*β*_*L*,_*β*_*R*_). We defined the intercepts as action pressure parameters as they determine the subject’s overall willingness to perform each action independent of sensory evidence.

In the models we used, the auditory stimulus variables were *A*_*L*_ and *A*_*R*_ for left and right auditory stimuli, each set to zero if the sound was on the other side, and to 1 if it was on its side or in the center. The visual stimulus variables were *V*_*L*_ and *V*_*R*_ for left and right visual stimuli, each set to zero if the visual stimulus was on the other side, and to *c*^*y*^ if the stimulus was on its side, where *c* is stimulus contrast and *γ* is an exponent to account for contrast nonlinearities in the visual system^2,10,14^

We compared 5 model formulations:

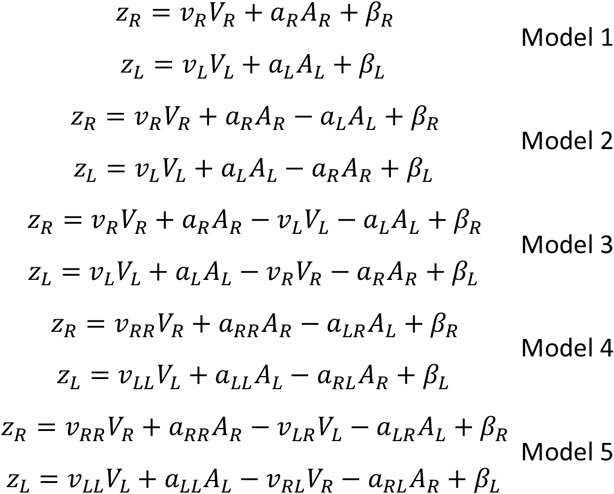

Model 1 assumed that stimuli only influence the decision variable for their respective side. Models 2 and 3 allowed for symmetric cross-influence (where a stimulus on one side can inhibit or promote the opposite choice). Models 4 and 5 allowed this symmetry to break, permitting stimulus-side-specific weights (e.g. a_LL_ or a_LR_) (**Supplementary Figure 1**).

To fit the models, we minimized the negative Log_2_Likelihood using *scipy*.*optimize*.*minimize*. We held out 20% of the trials to evaluate the model fits using the Log_10_Likelihood as a metric.

#### Optogenetic inactivation model

To evaluate the effects of optogenetic inactivation, we jointly fitted control and inactivation trials by expanding Model 3. We allowed each parameter to vary by the presence of inactivation (*I*) where *I* is an indicator variable (1 for inactivation trials, 0 for control):

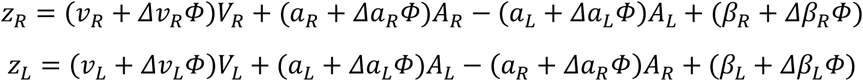

**Table 1.**
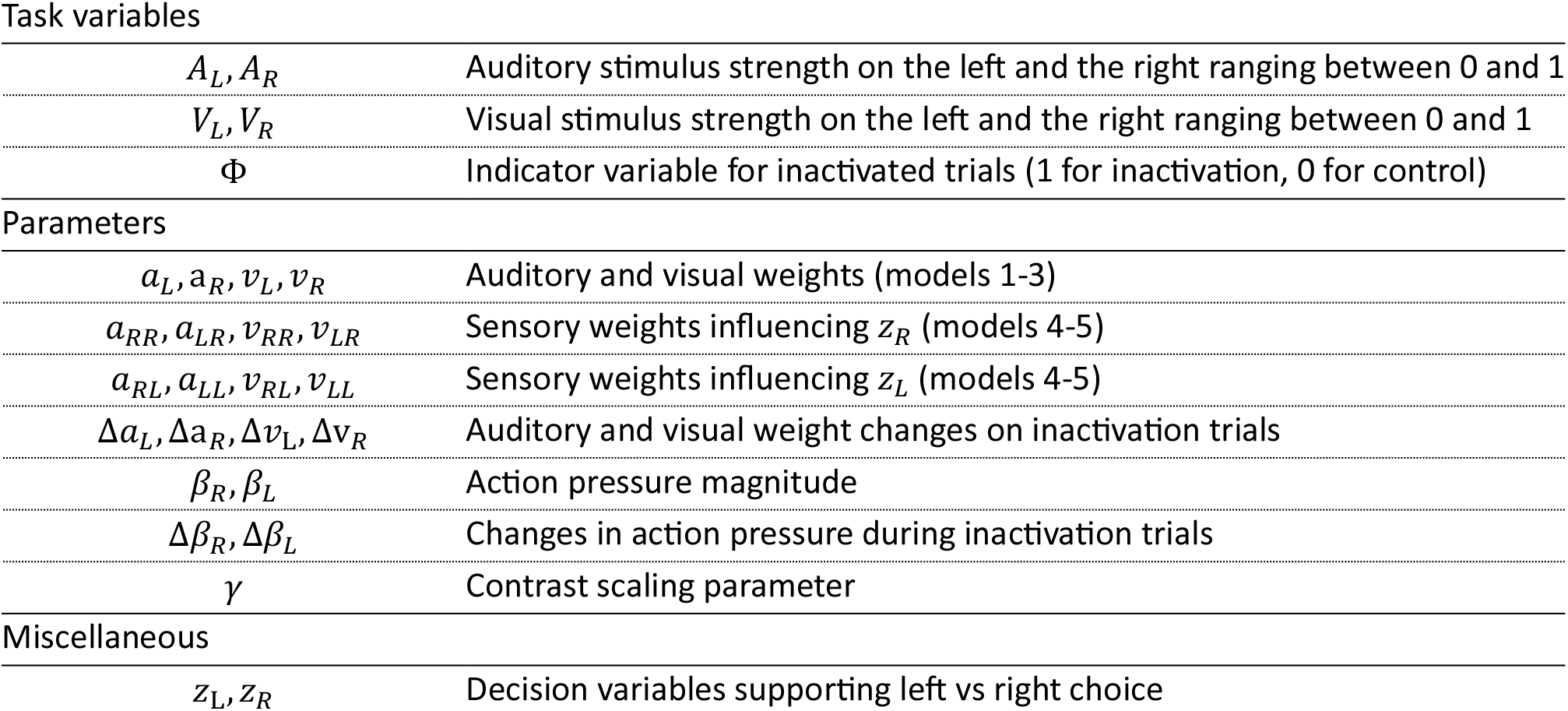
Summary notation of the 3-option model.

Task variables
|  |  |
| --- | --- |
| $A_L, A_R$ | Auditory stimulus strength on the left and the right ranging between 0 and 1 |
| $V_L, V_R$ | Visual stimulus strength on the left and the right ranging between 0 and 1 |
| $\Phi$ | Indicator variable for inactivated trials (1 for inactivation, 0 for control) |
| Parameters |  |
| $a_L, a_R, v_L, v_R$ | Auditory and visual weights (models 1-3) |
| $a_{RR}, a_{LR}, v_{RR}, v_{LR}$ | Sensory weights influencing $z_R$ (models 4-5) |
| $a_{RL}, a_{LL}, v_{RL}, v_{LL}$ | Sensory weights influencing $z_L$ (models 4-5) |
| $\Delta a_L, \Delta a_R, \Delta v_L, \Delta v_R$ | Auditory and visual weight changes on inactivation trials |
| $\beta_R, \beta_L$ | Action pressure magnitude |
| $\Delta \beta_R, \Delta \beta_L$ | Changes in action pressure during inactivation trials |
| $\gamma$ | Contrast scaling parameter |

**Table 1. Summary notation of the 3-option model.**
|  |  |
| --- | --- |
| $z_L, z_R$ | Decision variables supporting left vs right choice |

#### Inactivation effects as a function of LED powers

To quantify how bilateral optogenetic inactivation alters action pressure across continuous power combinations (*P*_*R*_ and *P*_*L*_ in mW/mm^2^) we first extracted trial-averaged empirical action pressure shifts (*Δβ*_*R*_, *Δβ*_*L*_) for each power condition by fitting a reduced 3-choice model where inactivation effects were restricted to intercept shifts:

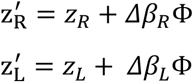

We then modeled the power-dependent structure of *Δβ*_*R*_ and *Δβ*_*L*_ according to Equation 6, where *I*_*L*_ and *I*_*R*_ represent the effective biological inactivation efficacy in the left and the right hemispheres generated by light delivery (which depends on NpHR3.0 load, and cannula location relative to the infected neurons):

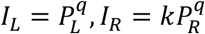

Here, *q* is a power sensitivity exponent nonlinearity of the effects of light stimulation and *k* denotes the efficacy of light stimulation in the right hemisphere relative to the left.

To evaluate model generalization, trials were split into two independent halves (train and test sets) to extract empirical Δ*β* parameters. The power model was fitted on the training split, and the out-of-sample prediction accuracy was quantified against the test set using the coefficient of determination (R^2^_score).

#### The audiovisual drift-diffusion model

To fit the DDM, we calculated reaction times by taking the timing difference between the first stimulus onset (auditory or visual) and the registered time that the mouse has brought the stimulus in the middle of the screen. We fitted the mouse choices and reaction times using a generalized drift-diffusion model that we implemented with the pyDDM package ^143^. The fullest model consisted of 16 parameters for testing bilateral inactivation and 15 parameters for bilateral inactivation.

**Table 2.**
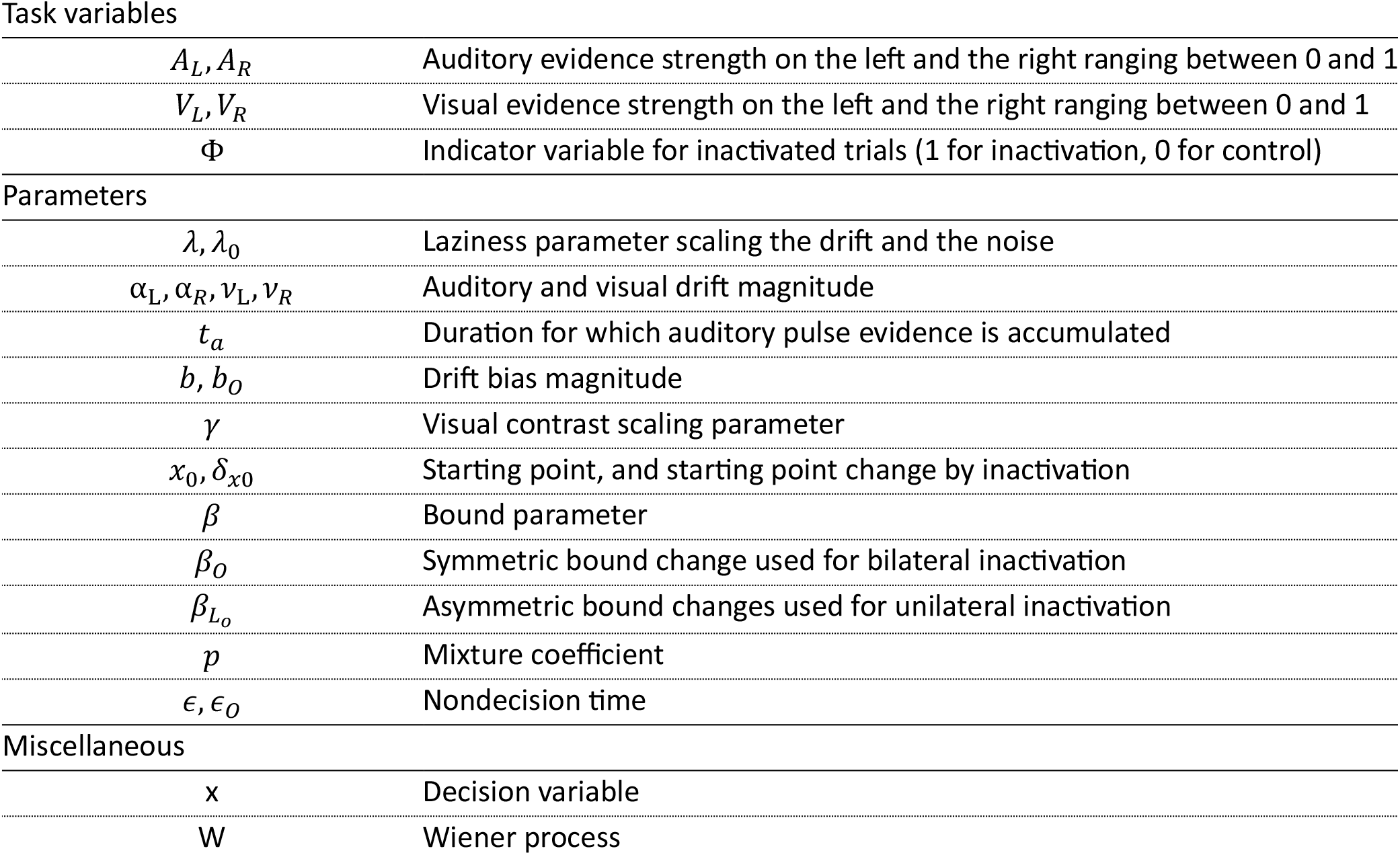
Summary of notation for describing the drift diffusion model.

Task variables
|  |  |
| --- | --- |
| $A_L, A_R$ | Auditory evidence strength on the left and the right ranging between 0 and 1 |
| $V_L, V_R$ | Visual evidence strength on the left and the right ranging between 0 and 1 |
| $\Phi$ | Indicator variable for inactivated trials (1 for inactivation, 0 for control) |
| Parameters |  |
| $\lambda, \lambda_0$ | Laziness parameter scaling the drift and the noise |
| $\alpha_L, \alpha_R, \nu_L, \nu_R$ | Auditory and visual drift magnitude |
| $t_a$ | Duration for which auditory pulse evidence is accumulated |
| $b, b_0$ | Drift bias magnitude |
| $\gamma$ | Visual contrast scaling parameter |
| $x_0, \delta_{x0}$ | Starting point, and starting point change by inactivation |
| $\beta$ | Bound parameter |
| $\beta_0$ | Symmetric bound change used for bilateral inactivation |
| $\beta_{L_0}$ | Asymmetric bound changes used for unilateral inactivation |
| $p$ | Mixture coefficient |
| $\epsilon, \epsilon_0$ | Nondecision time |

**Table 2. Summary of notation for describing the drift diffusion model.**
|  |  |
| --- | --- |
| $x$ | Decision variable |
| $W$ | Wiener process |

Momentary evidence x was modelled using a drift term µ, and a gain term *g*, such that

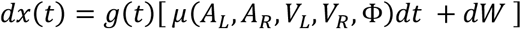

Here, drift is given at each timepoint t, from signed auditory and visual evidence and a constant bias (*b*) term, such that:

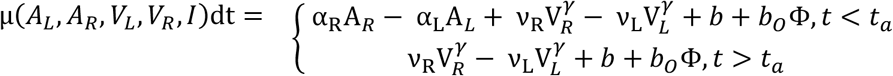

Introducing the gain term was necessary to fit NoGos and choices jointly (**Supplementary Figure 8**). The lazy gain was modelled as:

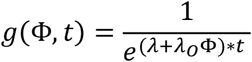

Here, *dx* was integrated from starting point of *χ*_*O*_(*I*), where until decision is reached determined by the function *f*_*bound*_(*I*). Trials that failed to reach either boundary within the decision window (*T* = 1.5 s) were classified as NoGo responses.

When testing the effects of bilateral inactivation we modelled the bound changes symmetrically, such that

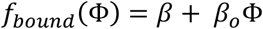

Here starting point was determined as a ratio between bounds:

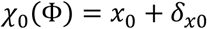

When testing the effects of unilateral SC inactivation, *χ*_*O*_(*I*) and *f*_*bound*_(*I*) were parameterized jointly, allowing asymmetric inactivation effects on the bound, such that the distance between the upper (ipsi choice) bound is fixed, and only the contralateral bound changes with inactivation^45^.

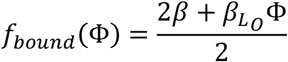

In this asymmetric model the bound to ipsilateral choices remains fixed, and thus effectively the zero position for the starting point is determined as:

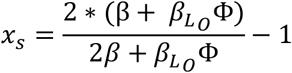

In this formulation we still allowed for a non-zero starting point and thus the effective starting point was determined as

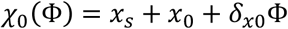

A decision was terminated when |*x*(*t*)| ≥ *f*_*bound*_(Φ).

All model’s RT distribution was given as a mixture between the first passage time distribution across the decision boundary and a uniform lapse distribution, with the mixture proportion determined by a fixed parameter *p*. The lapse component is uniformly distributed over the model’s time domain and contributes equally to left/right choice alternatives.

We also include a nondecision time given as

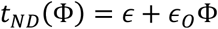

Therefore, the model’s predicted reaction time distribution is for leftward and rightward choices are given as

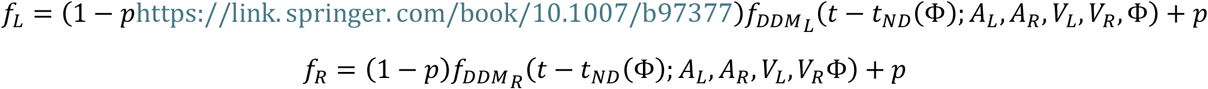

And the fraction of NoGos are given by the survival probability:

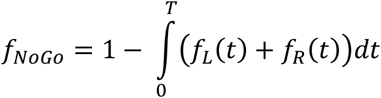

Trials were split into training and test-set using the *StratifiedShuffleSplit* method in *scikit-learn* to represent all trial types equally in the training and test set. Models were fit by minimizing the −Log_10_Likelihood, computed sum of the log first-passage time density evaluated at the observed RT for Left and Right trials, and the log survival probability for NoGo trials. This is the right-censored survival likelihood ^144^. Models were compared against each other using the same metric on the test set.

In this framework, we sought to assess the changes caused by SC inactivation of the SC by allowing additive changes to each term on inactivation trials. We found that some inactivation parameters were unnecessary (**Supplementary Figure 9**).

For the final model of bilateral inactivation, we fixed the inactivation parameters *ϵ* = 0, *λ*_*O*_ = 0, *δ*_*xO*_ = 0, *b* = 0, simplifying the model to 11 control and 1 inactivation parameter, *β*_*o*_.

For the final model of unilateral inactivation, we fixed *ϵ* = 0, *λ*_*O*_ = 0, *δ*_*xO*_ = 0, simplifying the model to 11 control and 2 inactivation parameters, *β*_*Lo*_& *b*_*o*_.

### Neural data analysis

#### Preprocessing

All recordings were sorted with pyKilosort ^130^ using default settings and curated using Bombcell ^131^ using custom thresholds selected after visually inspecting the data (minimum spike number=1500, minimum presence ratio = 0.7, maximum percentage of spikes missing = 20%, maximum refractory period violations = 0.5, **Supplementary Figure 2**c). Each shank’s anatomical alignment was corrected using electrophysiological landmarks (the silent zone between the cortex and the SC) using the IBL Ephys Atlas GUI^145^. This alignment was further validated by checking SCs visual receptive fields on each shank (**Supplementary Figure 2**a,b,d). For single unit analyses, we ensured unit independence by manually selecting one or two anatomically disparate recordings per mouse.

#### Mapping visual receptive fields from responses to sparse noise

We constructed a raster of white-square onset responses for each screen location by baseline-subtracting activity (200 ms window) and smoothing it with a 25 ms causal Gaussian kernel. For each square onset, we then computed the mean response over the 20–80 ms post-stimulus window, yielding a matrix of responses at every square location. We split the data into training and test sets (50% of the trials each). Using the training set, we fitted a two-dimensional gaussian tuning curve to estimate each unit’s receptive field. To identify units with a significant visual receptive field, we required that the fitted model explain more than 5% of the response variance on the test set.

#### Kernel fitting to passive and task responses

We modelled the firing rates *R*(*t*) of each individual neuron as a linear combination of event-related kernels, capturing responses to visual and auditory stimuli, task engagement and movement execution. We fitted neural activity both during task engagement and passive reply simultaneously.

Spike trains were aligned to first stimulus onset (*t*_s_, defined as the first sense that was registered to turn on in the DAQ) binned in 10 ms bins and smoothed using a 25 ms causal Gaussian kernel. Firing rates were z-scored using the mean and the standard deviation of the baseline period across all trials, and noise units and units that did not exceed a baseline variance of 1 (spike/s)^2^ were discarded. We only plotted neurons that passed Bombcell quality metrics in our final analysis.

To model these responses simultaneously, we constructed a time-tagged design matrix (**Supplementary Figure 3**b) where each event type was represented with a series of binary indicator variables spanning the kernel’s support window. Thus, when the model was fitted to the neural data, the resulting coefficients directly represented the learned, time-resolved shape of each kernel function at 10 ms resolution.

The predicted firing rate 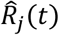 at time *t* relative stimulus onset during trial *j* was defined by the following linear model:

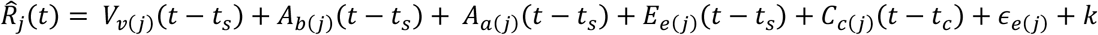

Where:

- *k* is a fixed baseline constant applied on all trials
- *t*_*s*_ and *t*_*C*_ indicate inset times of the stimulus and the choice action on trial *j*
- *Vv*_(_*j*_)_ represents one of the three for the three contralateral contrasts (low, medium, high) supported from 0 s to 0.45 s around stimulus onset (*t*_*s*_) on trials *j* when the visual stimulus was contralateral
- *A*_*b*_ is the auditory base kernel that is supported on trials where any auditory stimulus was presented (all trials except passive visual only trials), from 0 to 0.45 s around stimulus onset (*t*_*s*_)
- *A*_*a(j*)_ is the spatial auditory modulation kernel, representing either the ipsilateral (*A*_*ipsi*_) or the contralateral sound position relative to the neuron, supported from 0 s to 0.45 s around stimulus onset (*t*_*s*_). Applied essentially when sound was not at the center.
- *E* is the engagement gain kernel that allows could modulate stimulus responses when mice engaged in the task from 0 to 0.2 s around stimulus onset (*t*_*s*_). Applied on trials when mouse was making Left or Right choices
- *ϵ* is the engagement baseline kernel that accounts for constant baseline offset when mice are engaged in the task, supported throughout the entire trial *j*, applied on all trials when mice made Left, Right or NoGo choices.
- *C*_*C(j)*_ represents choice initiating action kernels, mapping to either ipsilateral or contralateral actions relative to the neuron, supported form −0.15 s to 0.1 s relative to choice action initiation *t*_*C*_

Firing rates were jointly fitted across all trial types including left, right, and NoGo choices and passive replay trials. For each trial, we fitted the firing rate from 0.4 s before stimulus onset until 0.1 s after the action initiation or, for trials without movement (NoGo & passive), 0.55 s after stimulus onset. Timepoints at which none of the kernels were supported were excluded. Trials with reaction times shorter than 50 ms were discarded to prevent stimulus kernels from explaining early movement-related activity. Trials with reaction times greater than 0.55 s were also discarded because late stimulus responses were too rare to estimate reliably. Finally, we removed trials in which mice moved the wheel before initiating their choice movement.

The linear model was fitted using Ridge regression (scikit-learn). After evaluating different levels of regularization via cross-validation, we used *α* = 10. Model performance for each neuron on held-out data was quantified using the coefficient of determination (*r*^2^). The contribution of each kernel was measured by computing the variance-explained (VE) score on the residual obtained after subtracting the predictions of all other kernels.

To estimate the fraction of significant neurons across SC, we used a sum VE threshold of 0.5% for sets of kernels (visual low, mid and high for visual, auditory base, ipsi, and contra for auditory, engagement baseline and gain for engagement, and action ipsi and contra for actions). We treated mice, rather than neurons, as independent units by computing the fraction of significant neurons within each mouse per area (minimum 10 neurons per area). We reported the mean fraction across mice and used a Linear Mixed-Effects model (*Fraction* ~ *Region* + (1|*MouseID*)) to compare regions (SCs vs SCm), evaluating significance with a one-sided Wald test on the region coefficient.

To quantify the relationship between variance explained of pairs of kernels we used a mixed effects rank correlation. This, essentially, is a hierarchical Spearman correlation, that accounts for subject related variability. For this, we converted variance explained scores to global ranks, and fitted a linear mixed effects model, such that (*Rank*_*Y*_ ~ *Rank*_*x*_ + (1|*MouseID*)). The partial correlation coefficient (ρ) was calculated from the fixed-effect t-statistic: 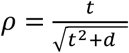.

#### Decoding choice

To decode choices, we computed the average firing rate of action and engagement neurons across time (from 200ms to 0 ms prior to action, or in 50ms bins to compute the temporal evolution of decoding accuracy). Action neurons were defined as >0.5% sum variance explained of the two action kernels, as previously, and engagement neurons were defined as >0.5% sum variance explained of the constant engagement and engagement gain kernels. Z-scored firing rates on each trial were used as predictors. In the extended model each neuron could acquire two independent weights to differentially contribute to left- and right decision variables according to Equation 5.

Notably, when fitting the model, *V*_*R*_ and *V*_*L*_ are scaled with *γ* as before, but one *γ* was obtained for each mouse, and was not fitted individually on experimental sessions.

We fitted this model on each recording session using the relaxed lasso approach ^146^. First, relevant neural features were selected using a multinomial logistic regression with an *L*_1_ penalty (scikit-learn) on the training set. Subsequently, the selected neural features and the stimulus predictors were fitted together without regularization by minimizing the negative log-likelihood.

In principle, this neural-augmented model can outperform a stimulus-only model in two ways: first, it can be more accurate, by better predicting choices on trials guided by internal or non-sensory processes (e.g., blank or conflict trials). Second, it can be more confident, by assigning higher odds to correctly predicted trials when neural predictors provide reliable evidence.

To understand these contributions, we assessed the models using the functional margin on the test set for each choice pair (Left/Right and Go/NoGo). For this, we calculated the predicted log-odds for each trial and multiplied them by the ground-truth outcome, encoded as (−1, 1). This transformation (typically used to fit SVM classifiers) aligns the classifier’s evidence with the actual behavior, such that positive values indicate a correct match between the neural signal and choice, while negative values indicate a mismatch (**Supplementary Figure 5**).

We decided to focus on highlighting model accuracy to focus on neural activity that captures internal processes we cannot capture by stimuli alone, such as attention, target selection or perceptual biases. To measure probabilistic accuracy, we computed the Brier score for choice pairs as:

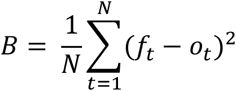

where *f* is the predicted probability of the choice an *o* is the actual outcome (1 if the choice occurred, 0 otherwise), for each trial *t*.

To test significant improvements within each region, we applied linear mixed effect models on the change in Brier score between the neural and the stimulus only models (Δ*B* ~1 + (1 | *MouseID*)), evaluating significance for a decrease with a one-sided Wald test on the intercept coefficient. To compare SCm vs MOs directly, we expanded the model to Δ*B* ~ *Region* + (1 | *MouseID*), evaluating significant Region contribution.

